# Stabilization of recurrent neural networks through divisive normalization

**DOI:** 10.1101/2025.05.16.654567

**Authors:** Flaviano Morone, Shivang Rawat, David J. Heeger, Stefano Martiniani

## Abstract

Stability is a fundamental requirement for both biological and engineered neural circuits, yet it is surprisingly difficult to guarantee in the presence of recurrent interactions. Standard linear dynamical models of recurrent networks are sensitive to the precise values of the synaptic weights, since stability requires all eigenvalues of the recurrent matrix to lie within the unit circle. Here we demonstrate, both theoretically and numerically, that an arbitrary recurrent neural network can remain stable even when its spectral radius exceeds 1, provided it incorporates divisive normalization, a dynamical neural operation that suppresses the responses of individual neurons. Sufficiently strong recurrent weights lead to instability, but the approach to the unstable phase is preceded by a regime of critical slowing down, a well-known early warning signal for loss of stability. Remarkably, the onset of critical slowing down coincides with the breakdown of normalization, which we predict analytically as a function of the synaptic strength and the magnitude of the external input. Our findings suggest that the widespread implementation of normalization across neural systems may derive not only from its computational role, but also to enhance dynamical stability.

**SIGNIFICANCE STATEMENT:** Neural circuits must remain stable to function correctly, yet strong recurrent connections, which are essential for complex computations, often hinder this stability. We demonstrate that divisive normalization, a canonical neural operation found across many sensory systems, can actively stabilize neural networks. By analyzing a biologically plausible model (ORGaNICs), we find that normalization allows circuits to remain stable even when recurrent interactions are strong enough to otherwise cause exploding neural dynamics. Furthermore, we identify a theoretical link between the breakdown of normalization and critical slowing down, a state where the brain recovers slowly from perturbations. This suggests that loss of normalization may serve as an early warning signal for the onset of pathological instabilities, such as seizures.

## II. INTRODUCTION

Divisive normalization is a form of multiplicative neuronal modulation occurring in the brain whereby the response of an individual neuron is divided by the summed activity of other similarly tuned neurons. Introduced in the 1990s to explain nonlinearities in the responses of neurons in the primary visual cortex [1, 2], it was later invoked to interpret a larger body of physiological data in the olfactory [3] and auditory [4] cortical areas, as well as in cognitive processes such as attention, working memory and value-based decision making [5–8]. Simply put, normalization explains how the response of a given neuron, which is selective for a specific stimulus, is suppressed by different stimuli that would elicit a weaker or no response if they were presented alone, as illustrated in Fig. 1a. Notwithstanding the general character of this neural computation, different biophysical mechanisms may perform normalization in different neural systems, including intracortical shunting inhibition [2, 9], thalamocortical synaptic depression [10], pre-synaptic inhibition [3], recurrent amplification (i.e., amplifying weak inputs more than strong inputs) [11–15], to name the most prominent ones. In our computational model, we implement divisive normalization via a multiplicative interaction between the principal neurons and a population of (secondary) inhibitory neurons, as illustrated in Fig. 1b,c. This model, which goes by the name of ORGaNICs [16], is much like the linear recurrent circuits introduced in the 1980s [17], but with a multiplicative gain on the recurrent term that implements normalization (see Fig. 1b,c).

**FIG. 1.**
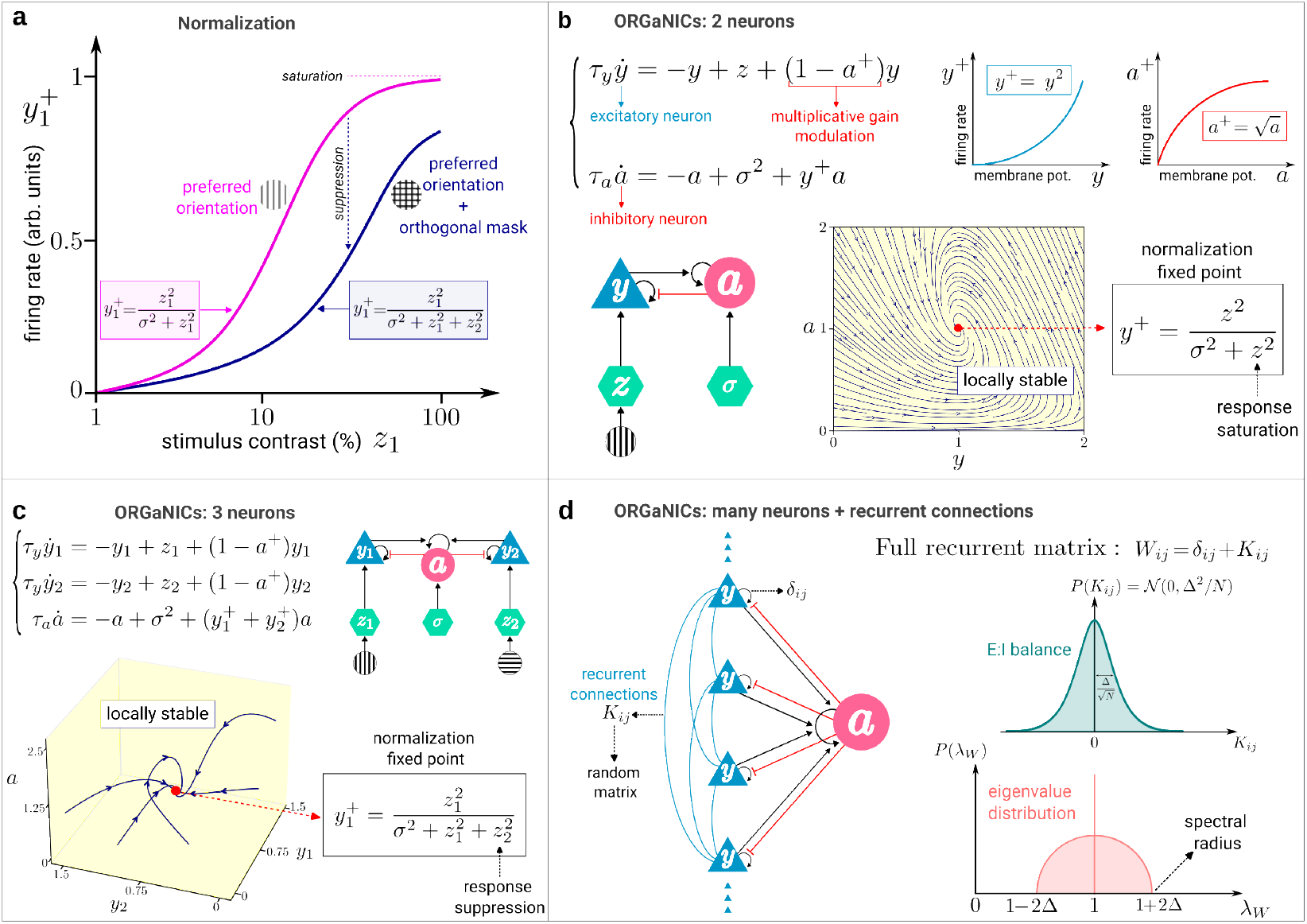
Normalization via ORGaNICs. **a**, An orientation selective principal neuron *y*_1_ in primary visual cortex (V1) fires when the stimulus orientation matches its preferred orientation (pink curve). The larger the stimulus contrast *z*_1_, the greater the strength of the response. The firing rate 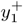 can be modeled by a nonlinear response function that saturates at high contrast. When a second grating stimulus *z*_2_ with orientation perpendicular to the preferred one (*viz*. an orthogonal mask) is presented simultaneously with stimulus *z*_1_, there is a rightward shift of the response function 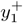 (blue curve). The suppressive effect of the orthogonal mask can be modeled by an extra term 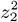 in the denominator of the response function. **b**, The saturation of the firing rate *y*^+^ at high contrast *z* can be obtained as the fixed point of the 2-neuron circuit in Eq. (2) involving the principal neuron *y* and a secondary inhibitory neuron *a* that acts on *y* as a multiplicative gain modulator. This fixed point is locally stable for any value of the time constants *τ*_*y*_, *τ*_*a*_ and the semisaturation constant *σ*. **c**, The suppressive effect of the orthogonal mask can be modeled by the fixed point of the 3-neuron circuit, where *y*_1_ and *y*_2_ respond selectively to the vertical and horizontal orientations, respectively, and *a* performs the multiplicative gain modulation on both *y*_1_ and *y*_2_. This fixed point is locally stable for any value of the parameters. Notice that neurons *y*_1_ and *y*_2_ do not interact directly, but only through neuron *a*, i.e., there are no recurrent connections between the principal neurons. **d**, Recurrent connections are included via the weight matrix *W* composed of the identity *I* plus a random matrix *K* modeling lateral synaptic connections between the principal neurons. The weights *K*_*ij*_ are E:I balanced (i.e., mean 0) and sampled from a symmetric Gaussian distribution such that the spectral radius of *W* is equal to 1 + 2Δ in the limit where the number of neurons *N* goes to infinity.

ORGaNICs have been proved to be unconditionally stable when the recurrent weight matrix is the identity [18]. Here we elaborate on the relationship between normalization and stability in the case of a generic recurrent weight matrix, such as a large random matrix drawn from the Gaussian Orthogonal Ensemble (GOE) [19]. We find that ORGaNICs models with recurrent weights drawn from the GOE ensemble are stable even when the spectral radius of the recurrent matrix is larger than 1, thanks to the normalization mechanism. Quantitatively, ORGaNICs push the stability limit of linear models by more than 100% (see phase diagram of stability in Fig. 5). Perhaps more importantly, we find that the transition to an unstable fixed point is preceded by critical slowing down [20, 21] in the neural dynamics (where the circuit is slow to reach the fixed point or to recover from small perturbations), the onset of which co-occurs with the breakdown of normalization in the neural responses. Remarkably, this result implies that the breakdown of normalization is an early warning signal for the loss of stability of the neural network, a signal we can predict analytically in terms of the recurrent interaction strength (i.e., the variance of the recurrent weights) and the magnitude of the external input.

## III. ORGaNICs MODEL OF NORMALIZATION

In its simplest form, normalization works by dividing a neuron’s total input by the sum of all inputs to *N* neurons in the normalization pool [1], expressed mathematically by the formula

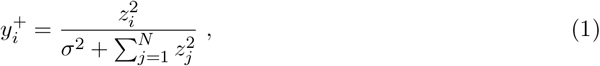

where 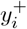 is the firing rate of neuron *i* defined as a nonlinear function of the membrane potential *y*_*i*_; *z*_*i*_ ∈ [0, 1] is its input drive-with 0/1 corresponding for example to 0*/*100% stimulus contrast-, defined as a weighted sum of the responses of a population of presynaptic neurons; and *σ* is the semisaturation constant, whose experimental value in primary visual cortex (V1) is *σ* ∼ 0.1 [22]. The sum in the denominator is the L2-norm of the input vector, which we assume to be bounded by 1 (this is always possible upon rescaling of the components *z*_*i*_). The purpose of the normalization mechanism is to normalize the output responses 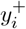 via the ratio between the input drive of an individual neuron and the input drives summed across all of the neurons [5, 16, 23–29]. Two important predictions of the normalization equation (1) as applied to visual cortex are illustrated in Fig. 1a, namely response saturation and cross-orientation suppression.

Since Eq. (1) describes a neural process that is static, it is natural to ask how the output responses 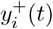 evolve in time towards the normalized state given by Eq. (1). That is: how does a neural circuit accomplish normalization? A mathematical way to achieve normalization dynamically is to couple the output responses of the principal neurons, 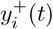, to a secondary neuronal population, represented by a single variable *a*(*t*), that acts as a multiplicative inhibitory modulator. The class of dynamical systems implementing divisive normalization in this way is known as ORGaNICs [16, 18]. The simplest ORGaNICs involve only two neurons and is described by the following dynamical equations

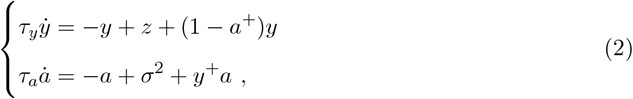

where *y*(*t*) and *a*(*t*) represent the membrane potentials (relative to an arbitrary threshold potential that we take to be 0) of the excitatory (E) and inhibitory (I) neurons, respectively, and *y*^+^ and *a*^+^ are the corresponding firing rates. The 2-neuron circuit described by Eq. (2) is depicted in Fig. 1b. The firing rate of the E neuron *y*^+^ is related to the membrane potential by squaring, *y*^+^ = *ky*^2^ [1, 30–33] (henceforth we set the dimensional proportionality factor *k* = 1), while the firing rate of the I neuron is given by 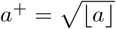, where ⌊*x*⌋ = max(0, *x*) (see Supplementary Section VII E and Fig. S10 for alternative activation functions); and *τ*_*y*_ *>* 0, *τ*_*a*_ *>* 0 are the neurons’ intrinsic time constants. The choice of multiplicative interactions in our model is justified by the extensive experimental literature demonstrating gain modulation in neural circuits and biophysical models of gain modulation [1–10, 34–38]. We posit that principal neurons and inhibitory neurons utilize distinct transfer functions using an approximately squaring function for principal cells and a compressive square-root function for the inhibitory modulator, as both convex [32, 33, 39] and concave [40] relationships are experimentally documented. The circuit in Eq. (2) models, for example, the response of a neuron in the primary visual cortex with *z* proportional to stimulus contrast, as seen in Fig. 1b. At the fixed point, the principal neuron *y* follows the normalization equation (1), i.e. 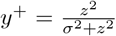, which explains the *saturation* of the firing rate at large contrast *z*. Moreover, this normalization fixed point is always locally stable [18] (see Fig. 1b, c).

Although it has been proved that a two-neuron ORGaNICs is unconditionally stable for any strength of the recurrent drive [18], the stability of a high-dimensional circuit with arbitrary recurrent connections has not been studied. Thus we ask: what happens when arbitrary recurrent connections (i.e. interactions) are included in the circuit? Do ORGaNICs still accomplish normalization? Is stability preserved?

To answer these questions we include recurrent connections between the principal neurons as described by the following set of differential equations

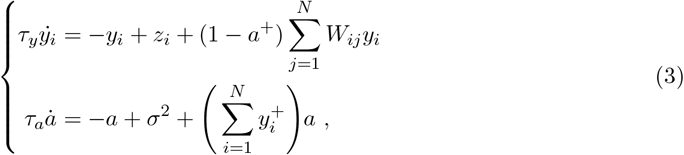

where the recurrent weight matrix *W* captures lateral connections between the principal neurons, as shown in Fig. 1d. Our goal is twofold: first, we determine the conditions on *W* and *z* such that normalization still approximately holds for the circuit in Eq. (3); second, we investigate the consequences of the breakdown of normalization, due to strong recurrent interactions, for the stability of the whole neural network.

## IV. LOSS OF NORMALIZATION AS AN EARLY WARNING SIGNAL OF NEURODYNAMICAL INSTABILITY

### A. Numerical solution of the fixed point

We start with a numerical study of the stability of the fixed-point of Eq. (3) and then we derive our analytical solution perturbatively, supported by the exact numerical result. We express the recurrent matrix as the sum of the identity plus a perturbation as

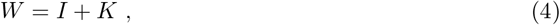

where *K* is a symmetric GOE random matrix [19] whose entries *K*_*ij*_ are independent and identically distributed Gaussian random variables with zero mean and variance Δ^2^*/N* if *i* = *j* or Δ^2^*/*2*N* if *i* ≠ *j*, corresponding to balanced excitation and inhibition. The scaling of the variance with 1*/N* ensures that the spectral radius *ρ*(*K*) does not grow with the number of neurons *N*, but is controlled only by Δ (specifically, *ρ*(*K*) = 2Δ, hence *ρ*(*W*) = 1 + 2Δ, see Fig. 1d). The choice of a random matrix to study the stability of large systems of differential equations can be traced back to the seminal work of May on the stability of complex ecosystems [41], that initiated a new field in theoretical ecology [42, 43] as well as the famous diversity-stability debate [44–46]. Methods based on random matrix theory are also well suited to model very large neural circuits whose experimental parametrization would be otherwise unfeasible [47]. The use of a random matrix to model recurrent interactions allows for conclusions that are generic and not specific to particular synaptic weights. We interpret negative weights as inhibition mediated by interneurons. Although this adds a transmission delay, we consider it negligible here. Here, we follow a similar approach with the goal of deriving a condition on the recurrent interaction strength Δ such that the output responses *y*_*i*_ still approximately satisfy the normalization equation (1), and then study the consequences of the breakdown of normalization on the system’s stability. We note, *en passant*, that a linear model (i.e. a model where *a* ≡ 0) would become unstable as soon as the spectral radius of *W* gets larger than 1, i.e. a soon as Δ *>* 0 (see Fig. 5). In contrast, the nonlinear model described by Eq. (3) can be stable even when *W* has spectral radius larger than 1, as we show next.

In Figure 2a,b we show the numerical solution of the fixed point of Eq. (3) (see Supplementary Section VII A for details on the numerical methods). We plot the mean and variance of the fixed point membrane potentials, *y*_*i*_, over the ensemble of random recurrent matrices *K*, as a function of the input drive *z* for Δ = 0.05 and Δ = 0.25. We find that the output responses, on average, still follow the normalization curve, i.e 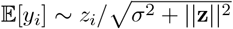 (noting that membrane potential in this model is the square root of firing rate), but pick up a variance that increases with increasing Δ. Note that the variance here is computed across realizations of the random connectivity (sample-to-sample variability) and should not be confused with the variance across presentations of the same stimulus (trial-to-trial variability [48]) that is not studied here since the the circuit is deterministic for a given instance of *W*. For sufficiently large Δ we observe that the circuit’s convergence to its fixed point, as measured by the real part of the largest eigenvalue *λ* of the Jacobian evaluated at the fixed point [49], becomes very slow (i.e. *λ* ∼ 0 corresponding to a convergence time 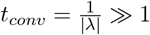, as seen in Fig. 2b. This phenomenon, called **critical slowing down**, is widely considered to be an important early warning signal that anticipates the system’s tipping point [20, 21, 50].

**FIG. 2.**
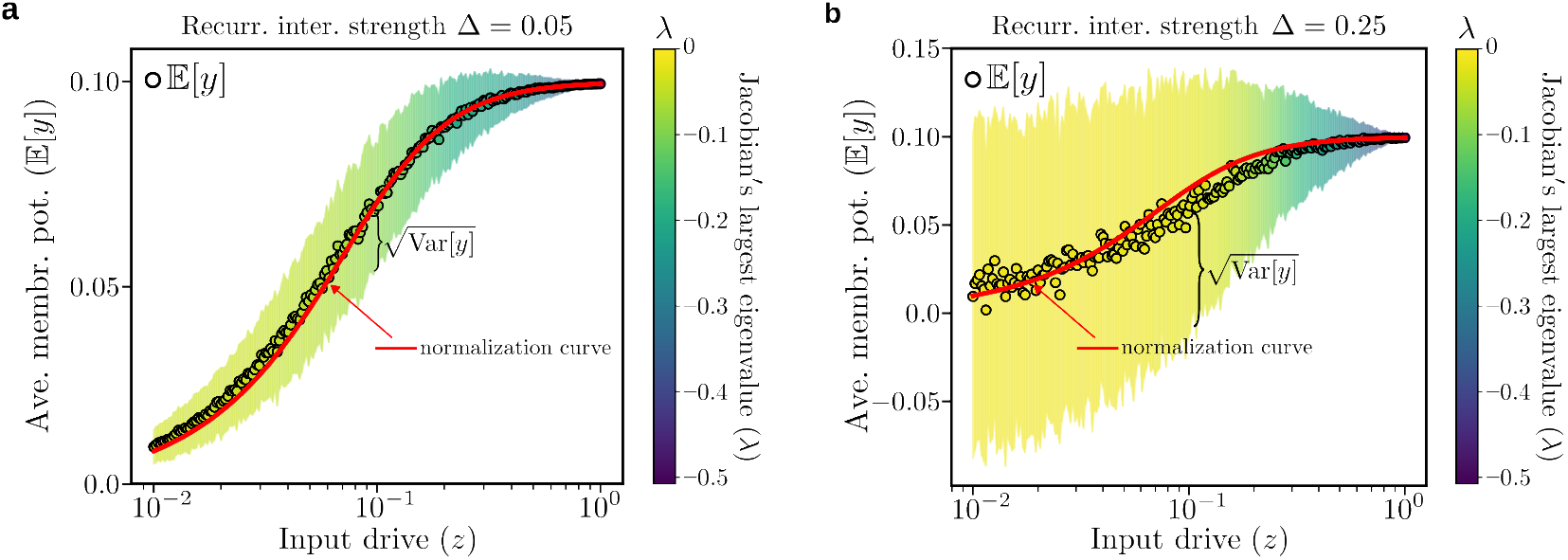
Numerical solution to the ORGaNICs’ fixed point equations. **a**, Fixed-point average membrane potential 𝔼 [*y*] (open circles) as a function of the input drive *z* (here we use *N* = 100 neurons and 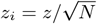, called *delocalized* input drive) for an E:I balanced recurrent network *K* with zero mean and std. dev. Δ = 0.05, obtained by solving numerically Eq. (3) using the explicit Euler method with time step *dt* = 0.05 *× τ*_*y*_. The semisaturation constant is *σ* = 0.1 and the neurons’ time constants *τ*_*y*_ and *τ*_*a*_ are equal. Each point is an average over 1000 realizations of the synaptic weight matrix *K*. The neural response still follows, on average, the normalization Eq. (1) (solid red curve), but picks up a variance across different random samples of recurrent synaptic weights, represented by the shaded area around the data points. The color code of the shaded area represents the real part of the largest eigenvalue of the Jacobian at the fixed point averaged over samples, whose value is well below 0 for all *z*. **b**, For Δ = 0.25 the average response is still normalized, but the variance (across random samples of the recurrent weights) is bigger than in (**a**) (see Fig. S1 for more Δ values). For sufficiently small input drives, the largest eigenvalue of the Jacobian at the fixed point becomes very small (*λ* ∼ 0). As a consequence, convergence to the fixed point occurs on time scales much longer than the time constant *τ*_*y*_ of individual neurons, a phenomenon known as critical slowing down (see also Fig. 3c).

To identify the onset of critical slowing down we plot in Fig. 3a the probability distribution *P* (*λ*) of the real part of the largest eigenvalue of the Jacobian at the fixed point for circuits with *N* = 1000 neurons, weak input drive *z* = 0.01, and different values of the recurrent interaction strength Δ. At small Δ, *P* (*λ*) has a gap from 0, which closes when Δ approaches the critical value Δ = Δ_*csd*_, signaling the onset of critical slowing down. To get a more precise estimate of Δ_*csd*_, we extrapolate the mean and variance of *P* (*λ*) in the limit *N* → ∞ via finite size analysis, yielding the asymptotic mean and variance shown in Fig. 3b (see Supplementary Section VII B and Figs. S2, S3 for details on the extrapolation to *N* → ∞). The variance goes to zero in the large *N* limit, meaning that *P* (*λ*) becomes a *δ*-function sharply peaked around its mean. The mean vanishes at Δ = Δ_*csd*_ and remains zero in the whole interval Δ_*csd*_ ≤ Δ ≤ Δ_*c*_ (Fig. 3b). In this interval the neural dynamics are very slow (compared to the neuron’s intrinsic time scale *τ*_*y*_) to reach the fixed point, as illustrated by some representative trajectories shown in Fig. 3c. Eventually, for Δ ≥ Δ_*c*_, the circuits enter first into limit cycles and then become unstable (see Fig. 5). The reason why the network remains stable even when the spectral radius of *W* is larger than one can be intuitively understood by noticing that the multiplicative term (1 − *a*^+^) in Eq. (3) dynamically scales the recurrent matrix and, in doing so, can effectively reduce its spectral radius.

**FIG. 3.**
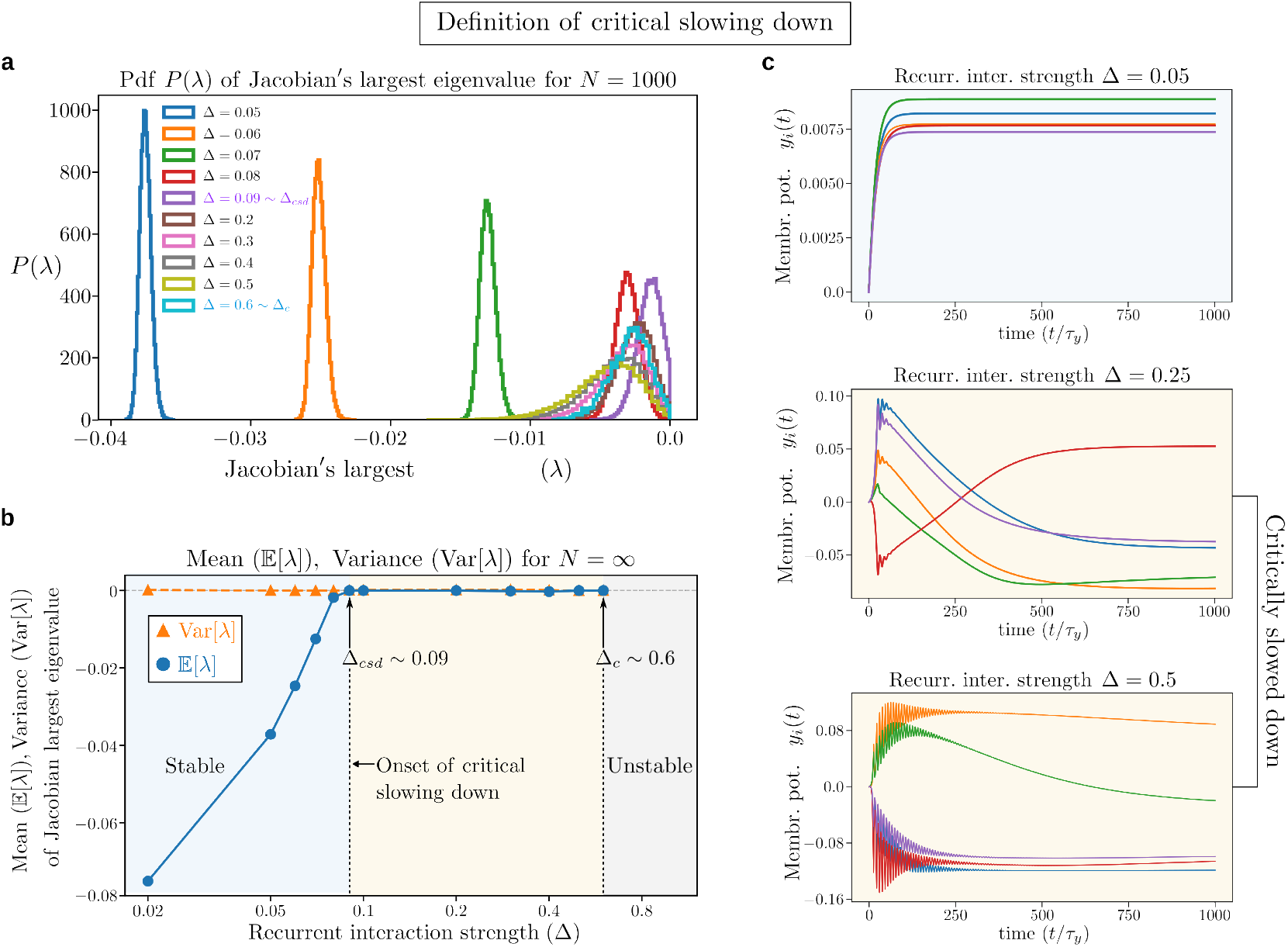
Definition of critical slowing down. **a**, Probability distribution of the largest eigenvalue of the Jacobian at the fixed point for a network with *N* = 1000 neurons (see Fig. S2 for different sizes); input drive 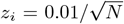 (very weak input drive); semisaturation constant *σ* = 0.1; and several values of the recurrent interaction strength Δ. For small Δ the distribution *P* (*λ*) is gapped from 0 and, as a consequence, the neural responses converge quickly to their fixed points. When Δ increases, the gap shrinks and then closes at Δ = Δ_*csd*_, signaling the onset of critical slowing down. For Δ_*csd*_ ≤ Δ ≤ Δ_*c*_ the neural responses converge slowly to their fixed points. For Δ ≥ Δ_*c*_ the neural circuits do not have stable fixed points, but exhibit limit cycles and, for even larger Δ, they eventually become unstable (see Fig. 5). **b**, Mean, 𝔼 [*λ*], and variance, Var(*λ*), of the largest eigenvalue of the Jacobian extrapolated to *N* → ∞ as a function of Δ (see Fig. S3 for details on the extrapolation). The model parameters’ values are as in (**a**). Since the variance is zero in the *N* → ∞ limit, *P* (*λ*) tends to a delta function *δ*(*λ* − 𝔼 [*λ*]), thus making the determination of Δ_*csd*_ well defined as the value at which the mean 𝔼 [*λ*] goes to zero. Slowing down persists up to the critical value Δ_*c*_, beyond which there are no stable fixed points (see Fig. 5). **c**, Representative trajectories of the neural responses *y*_*i*_(*t*) in the stable phase (Δ = 0.05) and in the critically slowed down phase (Δ = 0.25, 0.5), showing the slowness of the dynamics in reaching the fixed point. (see Fig. S4 for trajectories of all the neurons and Figs. S9, S12 for the analysis of the frequency of oscillations).

Next we demonstrate that the onset of critical slowing down occurs precisely when normalization of the neural responses breaks down.

### B. Loss of normalization predicts critical slowing down

To quantify the loss of normalization, we look at the mean and variance (across many instances of the recurrent matrix *K*) of the neural responses. As seen in Figure 2, the neural response *y*_*i*_ can be described by its expected value plus the standard deviation: the expected value follows the normalization equation and the standard deviation quantifies the departure from normalization. Therefore, neuron *i* loses normalization as soon as the standard deviation of its response is equal to its mean value, as given by the formula

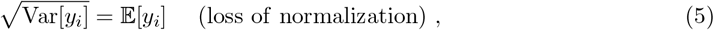

and illustrated in Fig. 4a,b. Equation (5) defines implicitly a threshold Δ_*loss*_(*z*) marking the boundary between the phase in which responses are normalized and the phase where they are not. We compare the loss of normalization threshold Δ_*loss*_(*z*) with the critical slowing down threshold Δ_*csd*_(*z*) in Fig. 5, showing excellent agreement between the two at all values of the input drive *z*, thus demonstrating that the onset of critical slowing down co-occurs with the loss of normalization of the neural responses, which represents our most important result.

**FIG. 4.**
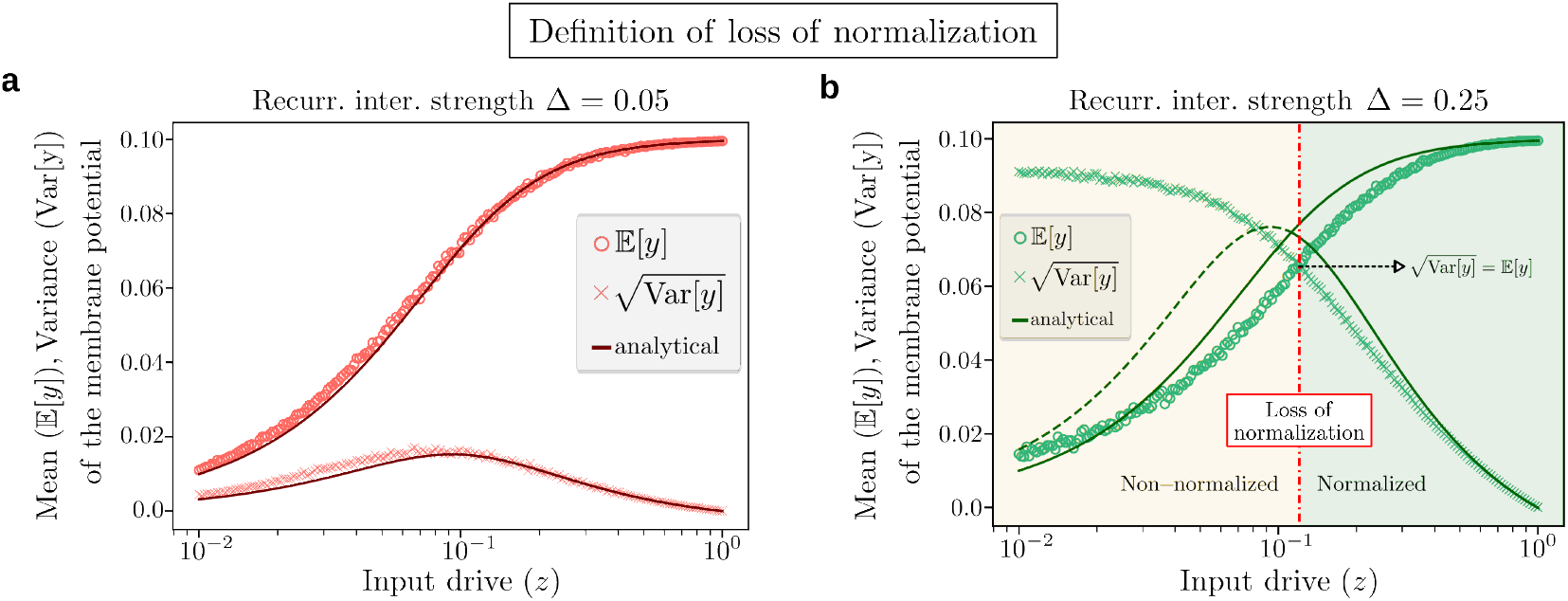
Definition of loss of normalization. **a**, Mean, 𝔼 [*y*], (circles) and standard deviation, 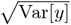, (crosses) of the fixed point membrane potential as a function of the norm *z* of the input drive 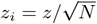 for an E:I balanced recurrent network *K* with zero mean and std. dev. Δ = 0.05. We used *N* = 100 neurons and averaged over 10^4^ realizations of *K*. The standard deviation is smaller than the mean for all values of *z*, so the neural responses are always normalized. The analytical approximations (solid curves) for the mean and standard deviation Eq. (7) of the response, computed with perturbation theory, show a good agreement with the exact numerical solutions. **b**, Same as in **a**, but using Δ = 0.25. The standard deviation is smaller than the mean 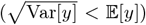 at large *z*, but it is larger than the mean 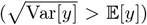 at small *z*. The value of *z* where the two curves cross each other, given by Eq. (5), defines the threshold at which the neural responses lose normalization (dashed red line). The analytical approximations are in good agreement with numerical simulations for almost all values of the input drive (notice the log scale on the abscissa), but become less accurate at small *z* where the responses are non-normalized and perturbation theory breaks down, another indication of a major shift in the circuit’s behavior.

**FIG. 5.**
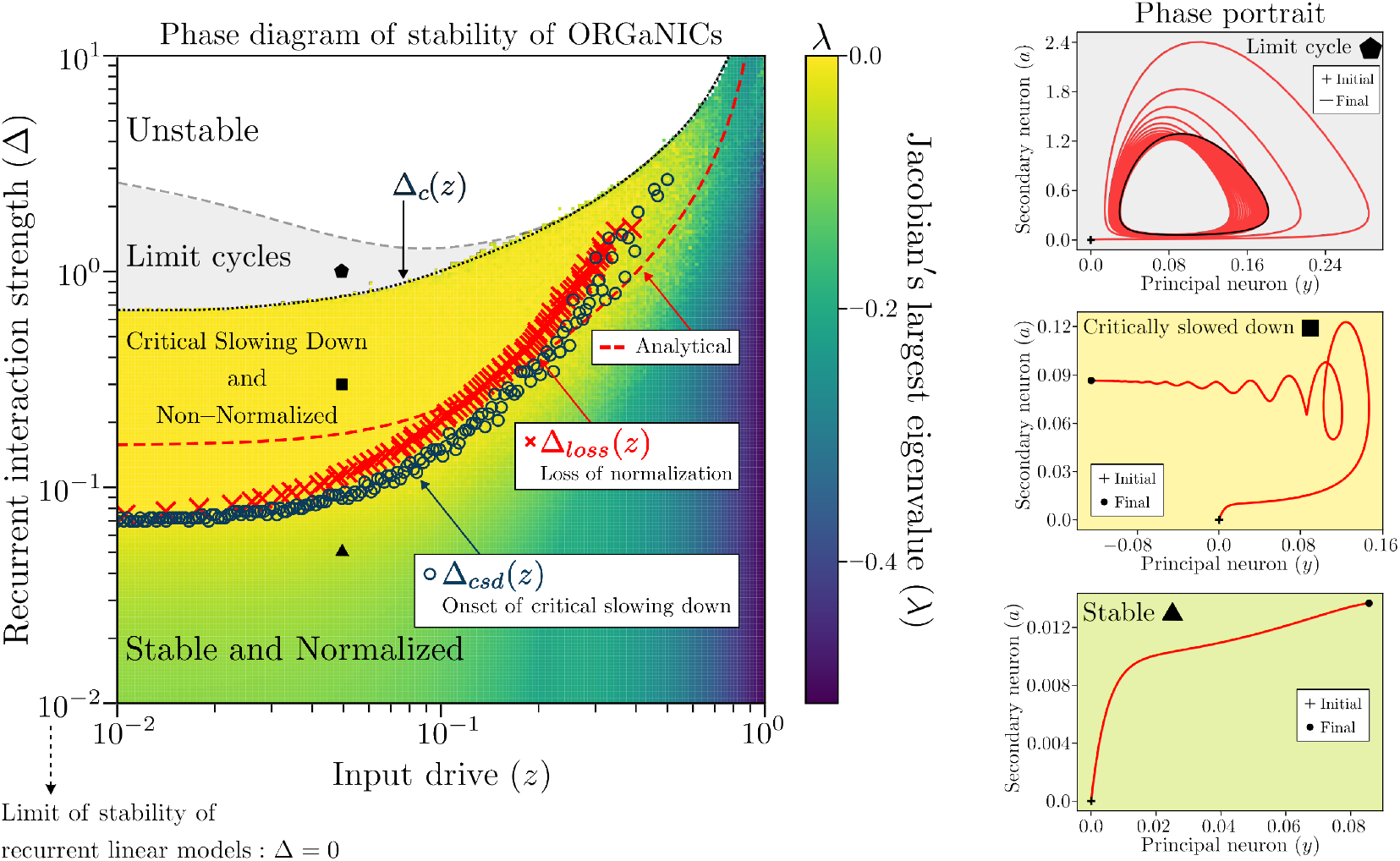
Loss of normalization predicts the onset of critical slowing down. Real part of the largest eigenvalue *λ* of the Jacobian at the fixed point in the (*z*, Δ) plane obtained by solving numerically Eq. (3) for E-I balanced networks with *N* = 100 neurons (see Supplementary Section VII F and Fig. S11 for the case of E-I imbalanced networks); delocalized input drive 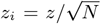; semisaturation constant *σ* = 0.1; neurons’ time constants *τ*_*y*_ = *τ*_*a*_; using a mesh of 200 *×* 200 values of *z* and Δ. Color represents the maximum value of *λ* across 100 random samples of the recurrent synaptic weights. Circuits with small Δ are stable at any value of the input drive *z* and converge quickly to their fixed point, as indicated by a strictly negative eigenvalue *λ <* 0 and by the *Stable* phase portrait in the (*y, a*) plane (where *a* is the inhibitory neuron). Conversely, for Δ_*csd*_ ≤ Δ *<* Δ_*c*_, the circuits exhibit critical slowing down, in that they approach the fixed point very slowly, as indicated by *λ* ∼ 0 and by the spiral attractor in the *Critically slowed down* phase portrait. The points marking the onset of critical slowing down (open blue circles) are determined by the closing of the gap in the distribution *P* (*λ*) (see Fig. 3), which define the curve Δ_*csd*_(*z*). The points defining Δ_*loss*_(*z*) (red crosses) represent the boundary between the normalized and non-normalized phases and are determined via Eq. (5) (see Fig. 4). Loss of normalization predicts well the onset of critical slowing down, i.e. Δ_*loss*_(*z*) ≈ Δ_*csd*_(*z*), thus providing a good early warning indicator of neurodynamical tipping points. For sufficiently large Δ the neural circuits exhibit limit cycles, as shown in the *Limit cycle* phase portrait, and for even larger Δ they become unstable. Notice, however, that simple linear recurrent models would become unstable as soon as Δ *>* 0, while adding normalization pushes the stability limit much further.

An immediate consequence of this correspondence is that we can predict theoretically the onset of critical slowing down by calculating Δ_*loss*_(*z*), which is a simpler quantity to estimate analytically, as explained next. To compute the mean and variance entering in equation (5), we must first find the fixed point of the dynamical system in Eq. (3), which, unfortunately, cannot be expressed in closed form. To overcome this obstacle, we use perturbation theory to approximate the exact solution. We look for a solution to the fixed point equations in the form of a series 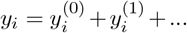, where 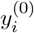 is given by the normalization equation (1), and 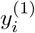 is of the same order of magnitude as the perturbation *K*. Inserting this expansion in Eq. (3) we find the approximate fixed point solution (see details in Supplementary Section VII C):

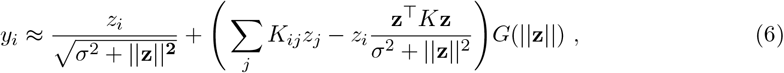

where 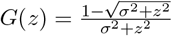 and ||**z**||^2^ = **z**^⊤^**z**. The last term on the right hand side of Eq. (6) quantifies the impact of the recurrent interactions on the normalization fixed point. Taking the expectation on both sides, and using the fact that the *K*_ij_ ‘s have zero mean, we find 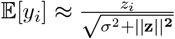, meaning that the neural responses still follow, on average, the normalization equation, as seen in Figure 4a,b. Departure from normalization is quantified by the variance of *y*_*i*_. The calculation of Var[*y*_*i*_] yields the following general expression (see Supplementary Section VII C for details)

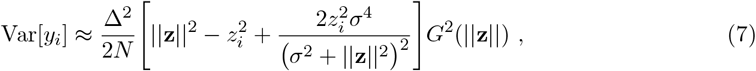

which depends on the magnitude ||**z**|| and shape *z*_*i*_ of the input drive. For example, we consider a *delocalized* input drive, i.e. 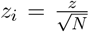, (the opposite case of a *localized* input drive is discussed in Supplementary Section VII C 2 and Figs S6, S7, S8, leading to qualitatively similar results) and find that Var 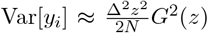, independent of *i*. In Figure 4a,b we plot the mean and variance of the neural response for two values of Δ, showing that the analytical approximations agree well with the exact numerical solution (see Fig. S5 for more values of Δ). Finally, by equating the mean and the standard deviation of the response, we find the threshold Δ_*loss*_(*z*) marking the boundary between the normalized and non-normalized phases as

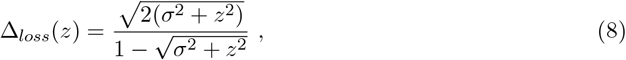

The analytical approximation given by Eq. (8) for the function Δ_*loss*_(*z*), shown in Fig. 5, is in good agreement with the exact numerical estimate. Since the discrepancy Δ_*csd*_ − Δ_*loss*_ is small relative to the magnitude of these thresholds both at low and hight input drive, i.e. Δ_*loss*_(*z*) ≈ Δ_*csd*_(*z*), then Eq. (8) can be used to predict the onset of critical slowing down in the neural dynamics from the magnitude of the *external* input and the strength of the *internal* recurrent weights. On the one hand, when *z* → 1 − *σ*^2^ ≈ 1 normalization is enganged robustly and the range of stability of the circuit extends indefinitely. On the other hand, when *z* → 0 the range of stability is narrowest.

It is important to note that loss of normalization is not the only possible cause of critical slowing down. We cannot exclude other mechanisms that might give rise to critical slowing down. Rather, we show: 1) that loss of normalization is correlated with critical slowing down, and 2) that critical slowing down can be used as an early warning signal for dynamical instability. Both of these theoretical results are experimentally-testable and potentially falsifiable.

## V. DISCUSSION

We have established, via numerical experiments and analytical calculation, that: *(i)* the nonlinear modulation of recurrent interactions via inhibitory neurons implementing divisive normalization makes neural networks more stable than unmodulated recurrent linear models; *(ii)* the breakdown of normalization, due to substantial recurrent amplification which is not compensated by an equally strong input drive, occurs concomitantly with the onset of critical slowing down in a broad class of random neural networks. Although loss of normalization could be detected, in principle, by recording changes in the recurrent weights, it may not be easily doable. A better way would be to measure the mean and variance of the firing rate across neuronal populations as a function of the input stimulus *z* and check when the mean-variance matching condition determined by Eq. (5) is fulfilled by tuning the stimulus level *z*. Critical slowing down, on the other hand, has been already measured experimentally and reported in the literature as a reliable biomarker for, e.g.. seizure forecasting [51].

Our results demonstrate that, at low input drives, increasing the recurrent synaptic strength turns the fixed point into a spiral attractor, as indicated by the damped oscillations in Fig. 3c and Fig. S9 (see Supplementary Section VII D for details on how to determine the frequency of oscillations). Crucially, the oscillations begin at the same parameter conditions where we observe the loss of normalization and onset of critical slowing down. This suggests a strong link between these two phenomena. Consequently, the detection of such recurrence-driven oscillations under a weak input drive could provide an experimental signature of a non-normalized neural circuit nearing a critical transition. Our computational framework predicts that increasing the recurrent interaction strength causes a failure of normalization, which in turn has been linked to diverse phenomena present in amblyopia [52, 53], epilepsy [54, 55], autism [56, 57], depression [58], and schizophrenia [59–62]. Furthermore, we predict that reduced normalization is accompanied by an increase in the effective time constant of the neuronal responses. This aligns with widespread experimental observations showing that when normalization is weaker (e.g., at lower stimulus contrast), effective time constants are larger [63, 64]. Consequently, we predict that circuits with impaired normalization will exhibit critical slowing down, which can be measured as the elapsed time to reach steady state. Increased neural variability, observed in patients with autism [65, 66], and noise correlations measured across trials of same stimulus presentation, are also characteristic markers of critical slowing down (see Supplementary Section VII G).

We noticed that, in the whole phase of critical slowing down, the spectrum of the Jacobian contains a large number of zero eigenvalues in the large *N* limit, corresponding to the emergence of multiple long time scales. Recently, it has been noted [67] that generating many long time scales in linear models requires fine tuning of the recurrent weights. In our model, many long time scales emerge for a broad range of values of the recurrent interaction strength, hence without fine tuning, suggesting that normalization (or other similar forms of multiplicative inhibitory modulation) might be the key mechanism to generate a full spectrum of slow modes in brain dynamics. A comprehensive analysis of the Jacobian’s spectrum, including the determination of the volume of zero modes, and the nature of the degenerate attractor will be presented elsewhere.

Finally, we distinguish our framework from Stabilized Supralinear Networks (SSNs) [68], which are cortical circuits with supralinear activation functions that achieve divisive normalization only approximately. Unlike ORGaNICs, SSNs do not exhibit the strict correlation between stability and normalization observed here, as they are inherently stable at weak inputs, where the normalization of responses is the lowest.

## VI. MATERIALS AND METHODS

### Model Simulation

We simulated the dynamics of Recurrent Gated Neural Integrator Circuits (ORGANICs) using networks of *N* = 100 up to *N* = 1000 principal neurons. The dynamics were integrated using the explicit Euler method with a time step *dt* = 0.05*τ*_*y*_, where *τ*_*y*_ = *τ*_*a*_ = 2*ms* is the membrane time constant. The recurrent weight matrix *W* was constructed as *W* = *I* + *K*, where *K* is a random matrix drawn from the Gaussian Orthogonal Ensemble (GOE). This formulation allows us to control the spectral radius of the recurrent interactions via a single parameter (the standard deviation of the weights), independent of network size.

### Stability Analysis

We determined the network stability by analyzing the Jacobian matrix of the system at its fixed point. The system was classified as stable if the real part of the largest eigen-value of the Jacobian, *λ*_*max*_, was strictly negative. Critical slowing down was identified when *λ*_*max*_ approached zero. We quantified the loss of normalization by comparing the standard deviation of the neural responses across random realizations of the weight matrix, against their mean response. Normalization is lost when the standard deviation equals or exceeds the mean 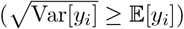.

## Code availability

The source code to perform all the calculations and plot the figures is available at https://github.com/shivangrawat/perturbed_organics.

## Acknowledgments

This work was supported by the National Eye Institute (R01-EY035242) and the National Institute for Mental Health (R01-MH137669). S.M. acknowledges the Simons Center for Computational Physical Chemistry (Simons Foundation grant 839534). This work was supported in part through the NYU IT High Performance Computing resources, services, and staff expertise. We thank J.B. Listman for discussions.

## Author Contributions

F.M. and S.R. contributed equally to this work.

## Additional information

Supplementary Methods accompany this paper.

## VII. SUPPLEMENTARY METHODS

### A. Numerical study of ORGaNICs’ fixed-point

To produce Fig. 2 in the main text we simulated an ORGaNICs network comprising *N* = 100 principal neurons with parameters set to *σ* = 0.1 and *τ*_*y*_ = *τ*_*a*_. For each chosen value of the recurrent interaction strength Δ, we generated an ensemble of 10^4^ recurrent connectivity matrices *W* = *I*+*K*. This was achieved by first sampling the entries of an auxiliary matrix *L* from a Gaussian distribution 𝒩 (0, Δ^2^*/N*), and then defining the symmetric interaction matrix *K* = (*L* + *L*^⊤^)*/*2. This prescription yields a symmetric random matrix *K*, whose entries are normally distributed according to

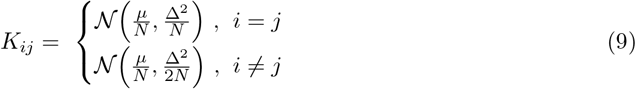

The network dynamics were simulated using the explicit Euler method (starting with a zero initial condition for all the neurons) with time step *dt* = 0.05 × *τ*_*y*_, using a delocalized input drive **z** where each component 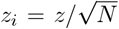 (ensuring ||**z**|| = *z*). We analyzed the steady-state behavior, identifying whether trajectories converged to a stable fixed point, diverged (indicating an unstable fixed point), or entered a limit cycle. For instances resulting in a stable fixed point, we computed the trial-averaged mean response 𝔼 [*y*_*i*_] and its standard deviation 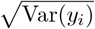 across the ensemble. Furthermore, we calculated the Jacobian matrix of the dynamical system at each stable fixed point using automatic differentiation [49]. Finally, as shown in Fig. 2 and Fig. S1, we plotted the mean response and its standard deviation as a function of the input drive *z* for different values of Δ. These plots are colored based on the average real part of the largest eigenvalue (in units of 1*/τ*_*y*_) of the Jacobian matrix across trials, indicating the slowest mode of the dynamics near the fixed point.

**FIG. S1.**
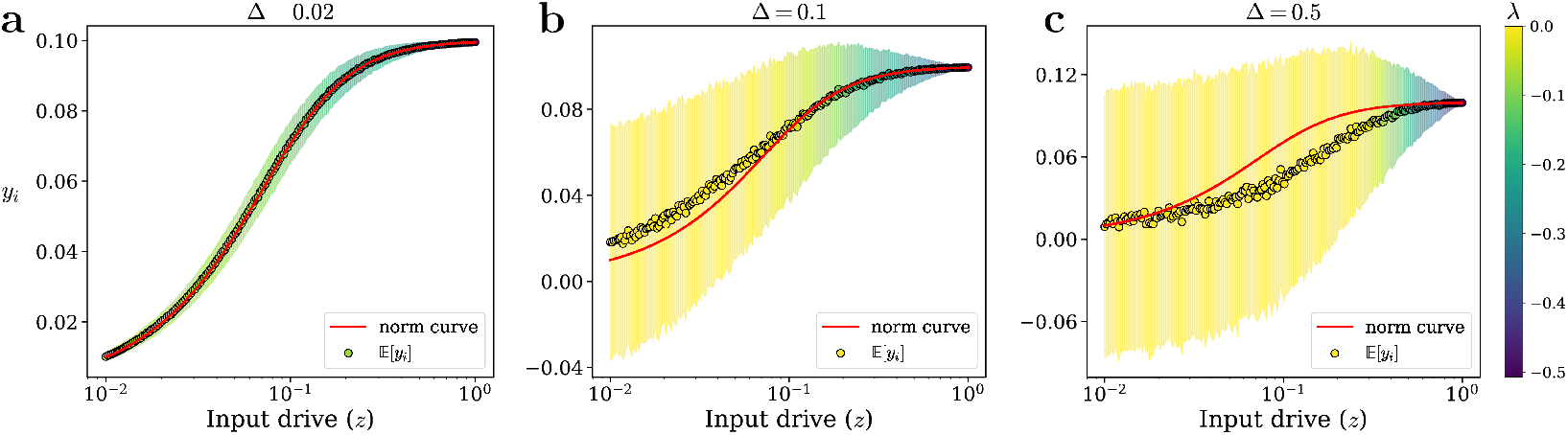
Numerical exploration of ORGaNICs’ fixed-point statistics. For each panel, we plot the fixed-point average response 𝔼[*y*] (dots) and its std. dev. (shaded area) as a function of the normalized input drive *z* (with 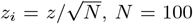), for an E:I balanced recurrence matrix *K* of zero mean and standard deviation Δ, with the same parameters as used in Fig. 2. The solid red curve indicates the normalization equation Eq.(1). The shading color encodes the real part of the largest Jacobian eigenvalue at the fixed point, averaged over samples (always *<* 0 when convergence is stable). **a**, Δ = 0.02: recurrent interactions are weak, yielding minimal variance around the normalization curve. **b**, Δ = 0.1: moderate recurrence induces variability in the responses across random samples of the recurrent weights at small *z*, but the mean follows the normalization curve. **c**, Δ = 0.5: strong recurrence dramatically increases the variability at small *z*; the mean also starts to deviate from the normalization curve.

### B. Finite size analysis of the distribution *P* (*λ*)

In this section, we investigate systematically the finite size behavior of the distribution of the largest eigenvalue of the Jacobian at the fixed point *P* (*λ*). In Fig. S2 we show *P* (*λ*) for several values of *N* and Δ. At fixed Δ, we find that *P* (*λ*) becomes sharply peaked as *N* increases and tends to a delta function, *P* (*λ*) → *δ*(*λ* − *λ*_*gap*_) in the limit *N* → ∞, where *λ*_*gap*_ is nonzero and negative when the circuit is stable (see Fig. S2a) and equal to zero when the circuit is critically slowed down (see Fig. S2b,c,d), i.e.

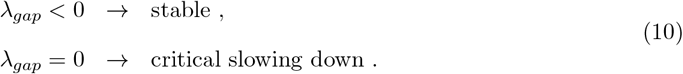

**FIG. S2.**
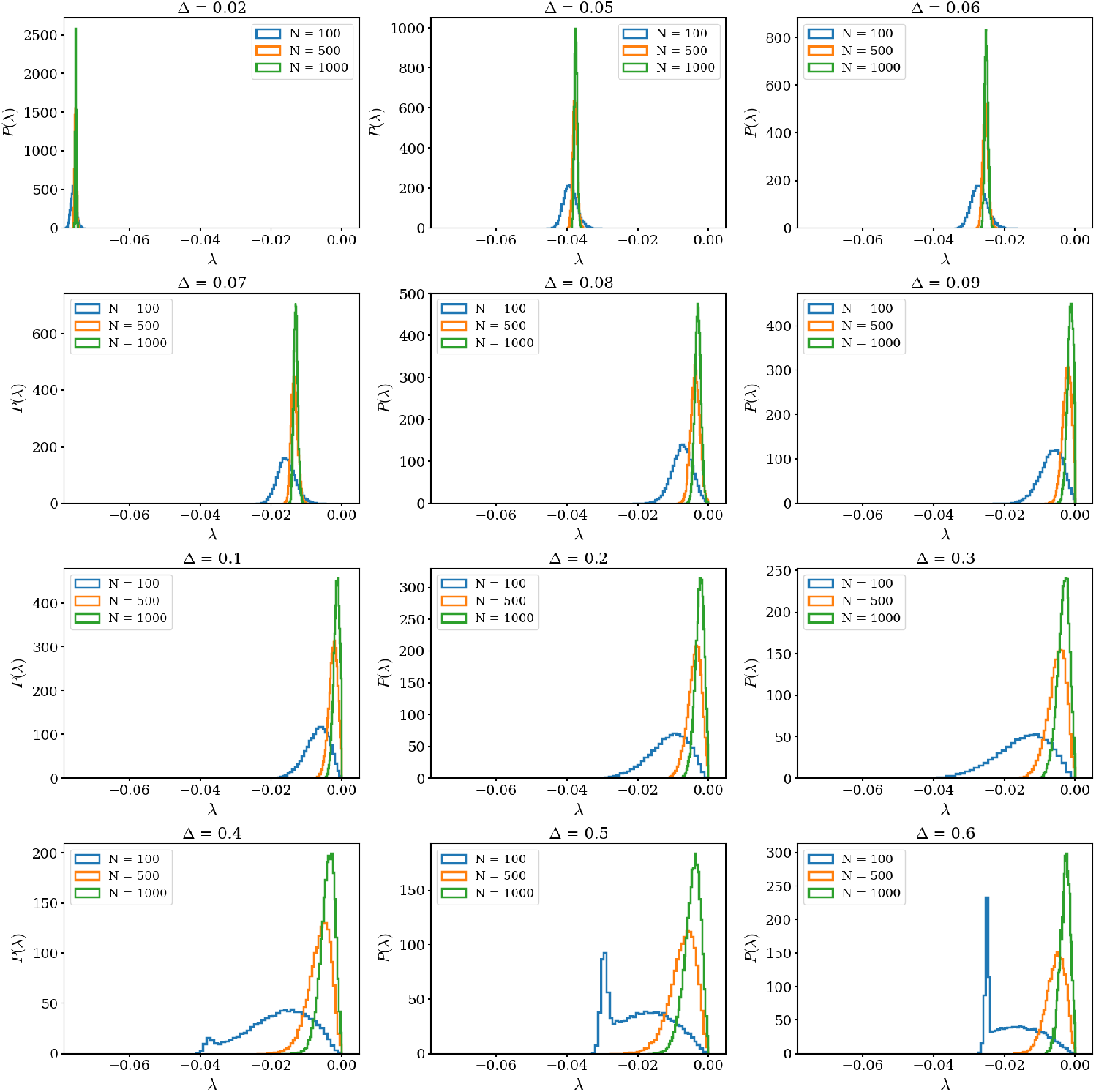
Distribution of the Jacobian’s largest eigenvalue (*λ*) for varying system sizes (*N*). Distribution of the largest eigenvalue of the Jacobian at the fixed point computed several values of Δ for different system sizes (*N* = 100, 500 and 1000). The input drive, simulation parameters, and the parameters of ORGaNICs are the same as those used for generating Fig. 3 in the main text. Each panel plots the distribution for different values of Δ. As system size increases, finite-size fluctuations narrow, sharpening the gap edge and more clearly revealing the approach of the rightmost eigenvalue toward zero at Δ ≈ 0.09. For Δ *<* Δ_*csd*_ (panel a), all sizes exhibit a clear gap from zero; for Δ ≥ Δ_*csd*_ the largest-*N* curve touches zero most sharply.

**FIG. S3.**
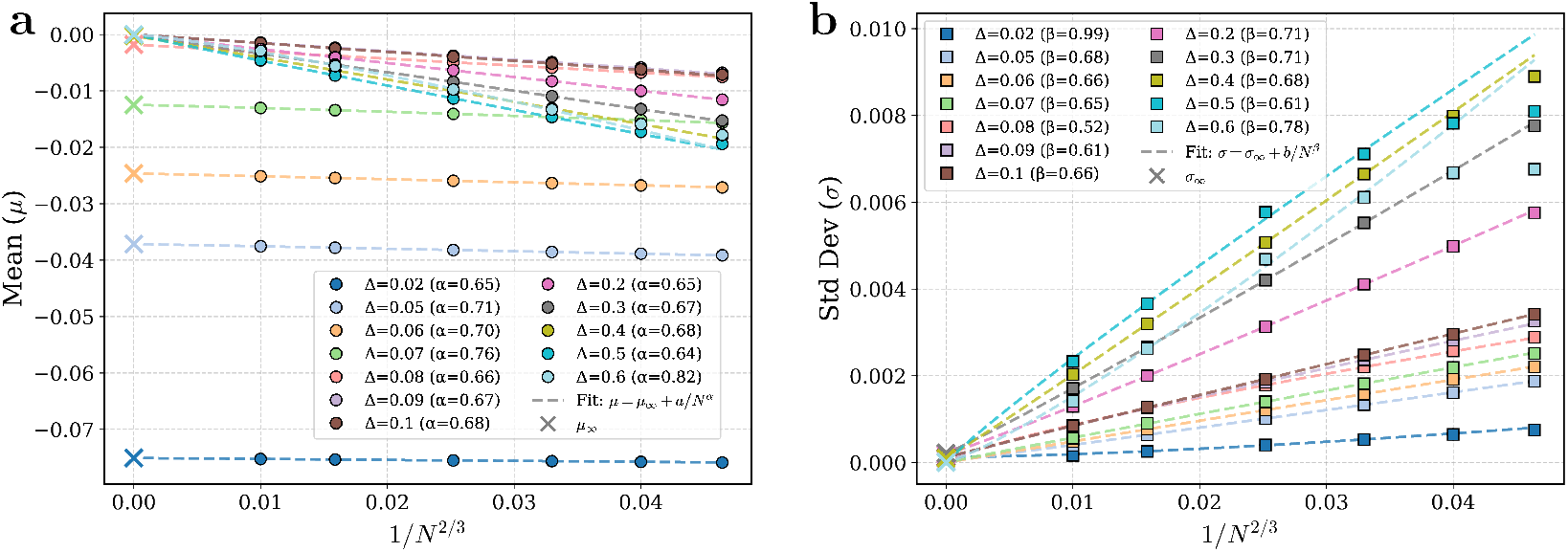
Finite size scaling analysis. The mean and standard deviation of the largest eigenvalue of the Jacobian are shown. The input drive, simulation parameters, and the parameters of ORGaNICs are the same as those used for generating Fig. 3 in the main text. **a**, The mean *µ* of the largest eigenvalue *λ* of the Jacobian matrix at the fixed point as a function of 1*/N* ^2*/*3^ for several values of Δ. Dashed lines represent fits following the functional form *µ* = *µ*_∞_ + *a/N*^*α*^, where *µ*_∞_, *a*, and *α* are fitting parameters and the fits are performed using the four largest system sizes. The ‘x’ markers correspond to the extrapolated mean for the infinite system size (*µ*_∞_). The values *α* in the legend correspond to the fitted slopes and they are close to 2*/*3 for nearly all values of Δ. *µ*_∞_ vanishes for all values of Δ where we observe critical slowing down (Δ ≳ 0.09). **b**, The standard deviation *σ* of the largest eigenvalue *λ* of the Jacobian matrix at the fixed point as a function of 1*/N* ^2*/*3^ for several values of Δ. Dashed lines represent fits following the functional form *σ* = *σ*_∞_ + *b/N*^*β*^, where *σ*_∞_, *b*, and *β* are fitting parameters, and fits are performed using the four largest system sizes. The fits extrapolate to a vanishing standard deviation for the infinite system size (*σ*_∞_ ≈ 0), indicating that fluctuations of *λ* vanish in the thermodynamic limit.

**FIG. S4.**
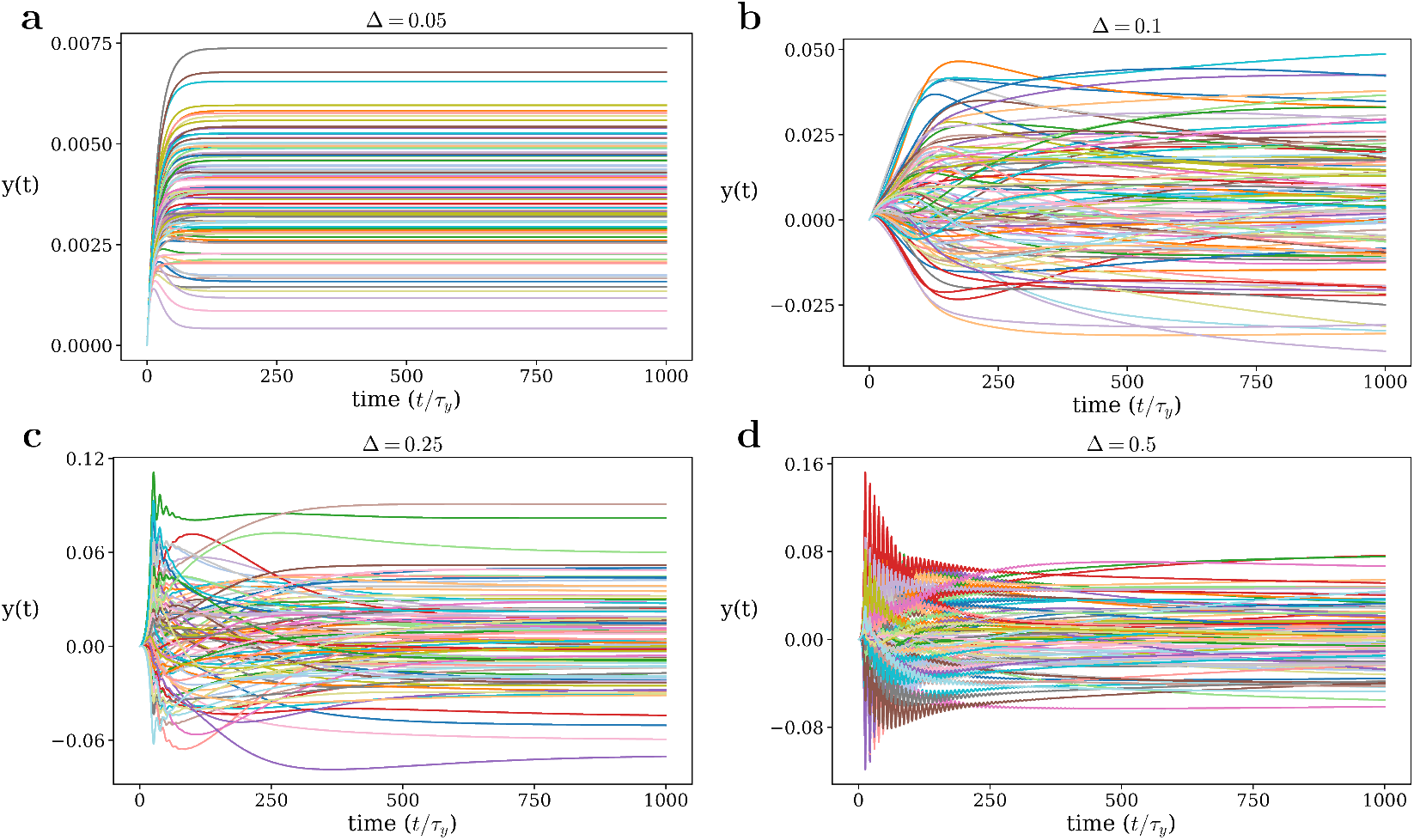
Neuronal trajectories for different recurrent synaptic strength Δ. Each curve traces the time evolution of a distinct principal neuron’s response for a given value of Δ (recurrent interaction strength). The input type and the parameters of ORGaNICs are the same as those used for generating Fig. 3 in the main text. We plot the trajectories for 100 neurons selected randomly from the 1000. In the stable regime (Δ = 0.05), trajectories converge rapidly, whereas in the critical-slowing regime (Δ = 0.10, 0.25, and 0.50) convergence is markedly slower.

### C. Analytical calculation of the threshold for loss of normalization

Let us consider the ORGaNICs fixed point equations obtained by setting to zero the time derivative in Eq. (3) of the main text:

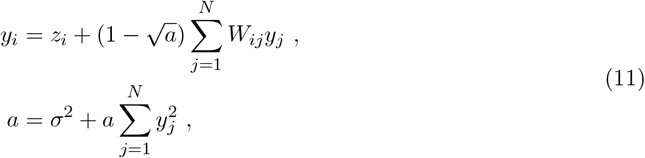

where we used the fact that firing rates are related to membrane potentials via

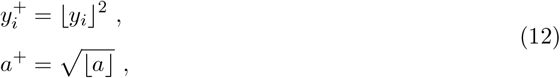

and we further assumed that *a* ≥ 0, which can be checked *a posteriori* to always hold true. To expand around the identity matrix we set

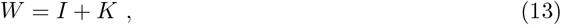

where *K* is a small correction. The conditions under which the perturbation *K* can be considered small with respect to the identity will be deduced later on in our calculation. Inserting Eq. (13) into Eq. (11) we obtain

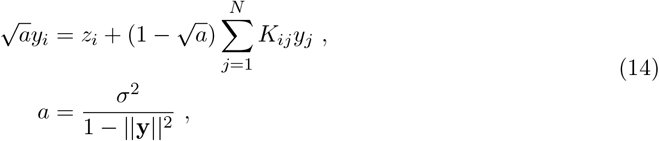

where we defined the squared norm as 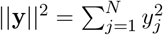. We look for a solution to Eq. (14) in the form of a series

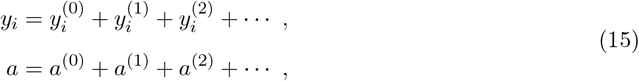

where 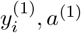 are of the same order of magnitude of the perturbation *K*, the quantities 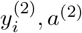 are of second order, and so on. To find the first approximation, we substitute 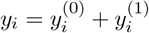 and *a* = *a*^(0)^ + *a*^(1)^ in Eq. (14) and we keep only terms up to the first order, thus obtaining

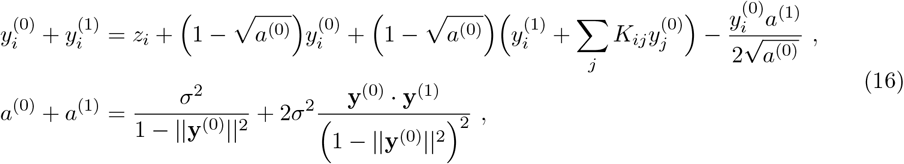

where 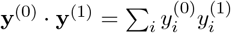 is the usual dot product. Equating the terms of order zero on both sides of Eq. (16) we obtain

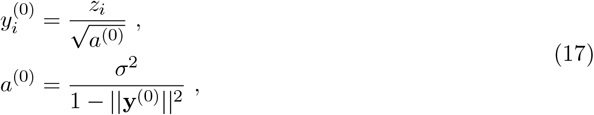

which, as it should, is equivalent to the normalization equation

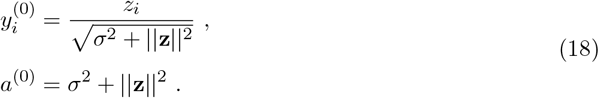

To find the first order corrections 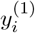 and *a*^(1)^ we equate the terms of order one on both sides of Eq. (16) and we get

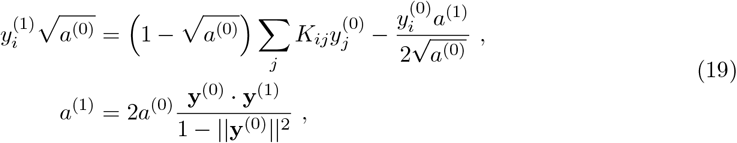

where in the equation for *a*^(1)^ we have used the definition of *a*^(0)^ given in Eq. (17). To solve Eq. (19) we multiply the first equation by 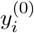 and, after summing over *i*, we find

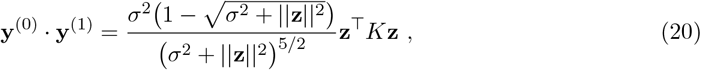

from which we can compute *a*^(1)^. Substituting this result into Eq. (19) we can express 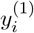 as a function of *z* and *K* as

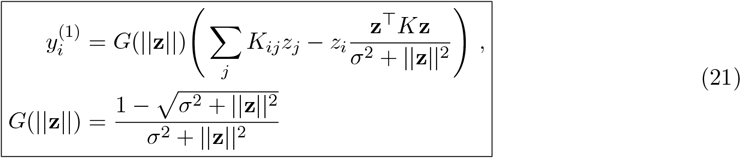

Having found the general form of the first order correction, we move next to consider the case of a random matrix *K* sampled from the so-called Gaussian Orthogonal Ensemble (GOE).

We consider the ensemble of symmetric random matrices *K*, whose entries are normally distributed according to

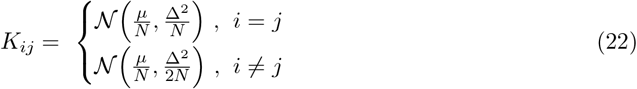

We can compute the average of 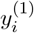 in Eq.(21) straightforwardly and find

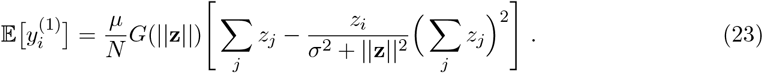

A little bit of algebra yields the following expression for the second moment

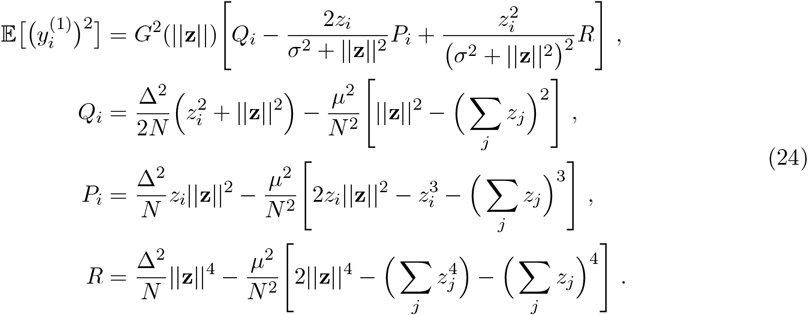

Having found the general expressions for the first and second moments of 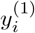, next we discuss the case *µ* = 0 (E:I balance), corresponding to having an equal number (on average) of positive and negative synaptic weights. Mathematically, this is obtained by setting to zero the mean (*µ* = 0) of the random matrix entries *K*_*ij*_. The mean and variance of the perturbation 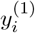 simplify considerably and read

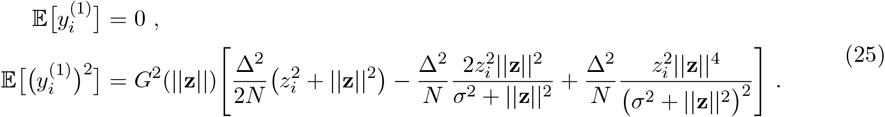

In the following, we will consider two types of input drives, a **delocalized** input drive, characterized by a vector **z** with all entries *z*_*i*_ equal to

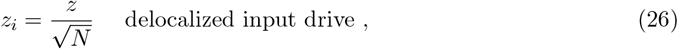

and the case of a **localized** input drive where all entries are equal to zero but one, for example *z*_1_, and denoted

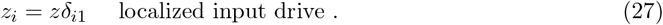

#### 1. Delocalized input drive

When the input drive is delocalized, the variance of the perturbation becomes

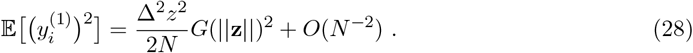

**FIG. S5.**
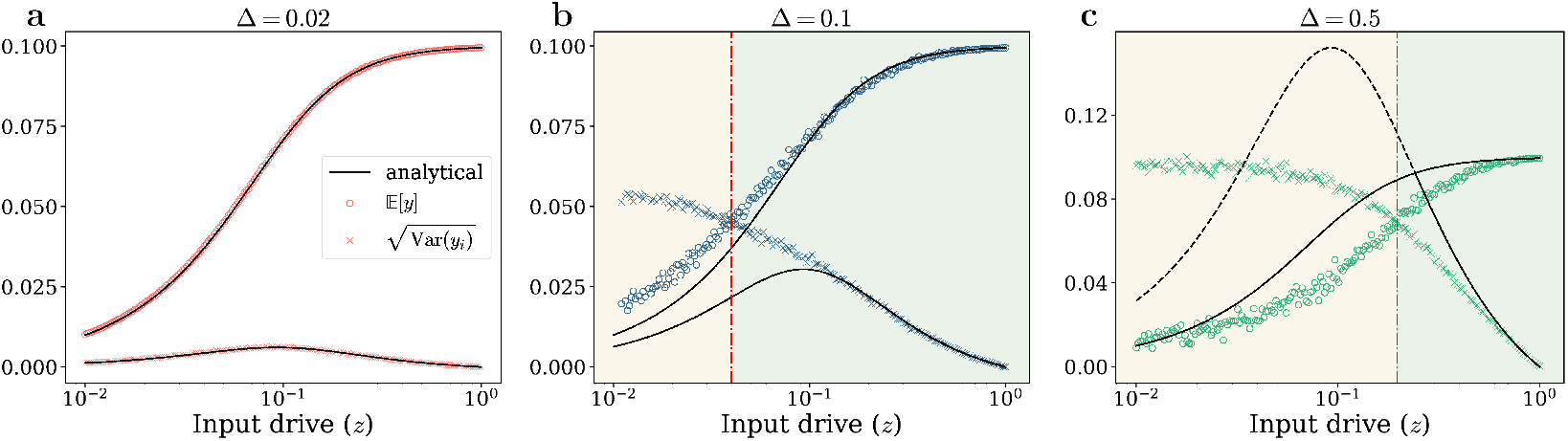
Loss of normalization. Following the analysis presented in Fig. 4, this figure illustrates the mean (circles) and standard deviation (crosses) of the fixed point neural responses versus the input norm *z* for three additional values of the recurrent interaction strength: Δ = 0.02, Δ = 0.1, and Δ = 0.5. The network consists of *N* = 100 neurons, and each point is found using 1000 realizations. **a**, Δ = 0.02, the std. dev. remains below the mean across all input drives, indicating preserved normalization. **b**, Δ = 0.1 and **c**, Δ = 0.5, the std. dev. exceeds the mean at small *z*, demonstrating loss of normalization, defined by the crossing point (dashed red line). The theoretical predictions from perturbation theory (black curves) match the numerical simulations well in the normalized regime (mean *>* std. dev.). Discrepancies increase at small *z* for larger Δ, where normalization is lost.

The threshold Δ_*loss*_(*z*) separating the phase where responses are normalized from the phase where they are not is obtained by equating the mean of the response 𝔼[*y*_*i*_] to its standard deviation 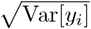, yielding

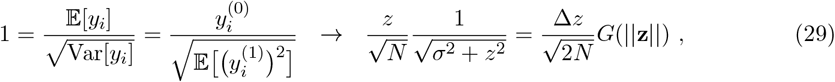

from which we obtain

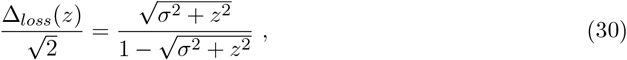

which is Eq. (8) in the main text. In Fig. S5 we compare the analytical approximations for the mean and variance of the responses with the exact numerical values. The agreement is excellent at small Δ, since the neural responses are always normalized, i.e. normalization always holds. For larger Δ the analytical and numerical results also agree well at large input drive, where the responses are normalized. At small *z*, normalization breaks down as well as the analytical approximation.

**FIG. S6.**
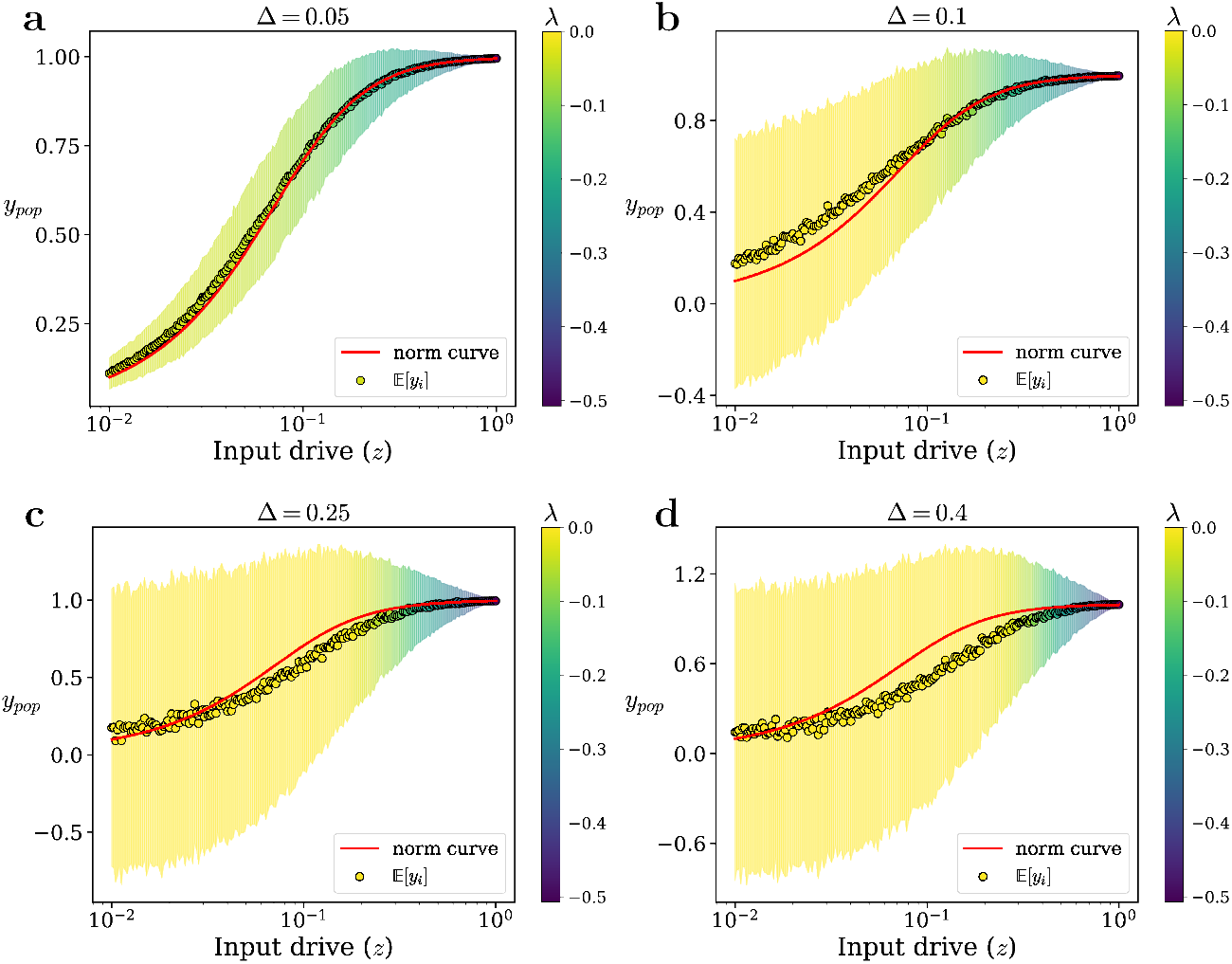
Numerical solution of the ORGaNICs’ fixed point equations for <u>localized input drive</u>. **a**, Fixed-point average response 𝔼[*y*_*pop*_], defined in Eq. (32), as a function of the input drive *z* (chosen as *z*_*i*_ = *zδ*_1*i*_) for an E:I balanced recurrence matrix *K* with zero mean and recurrent interaction strength Δ = 0.05 and *N* = 100 neurons, obtained by solving numerically Eq. (3) using the explicit Euler method with time step *dt* = 0.05 *× τ*_*y*_. The semisaturation constant is *σ* = 0.1 and the time constants *τ*_*y*_ = *τ*_*a*_. Each point is an average over 1000 realizations of the synaptic weights *K*_*ij*_. The neural responses still follow, on average, the normalization Eq. (1) (solid red curve), but pick up a variance in presence of recurrent connections, represented by the shaded area around the data points. The color code of the shaded area represents the real part of the largest eigenvalue of the Jacobian at the fixed point averaged over samples, whose value is well below 0 for all *z*. **b, c, d** For Δ = 0.1, 0.25, 0.4 the average response is still normalized, but the variance is bigger than in (**a**). For sufficiently small input drives, the largest eigenvalue of the Jacobian at the fixed point vanishes and, as a consequence, convergence to the fixed point occurs on long time scales, a phenomenon known as critical slowing down.

#### 2. Localized input drive

In this section, we show that our results and conclusions are the same for the localized input. We consider the extreme case where **z** is a one-hot vector with

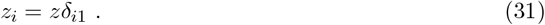

**FIG. S7.**
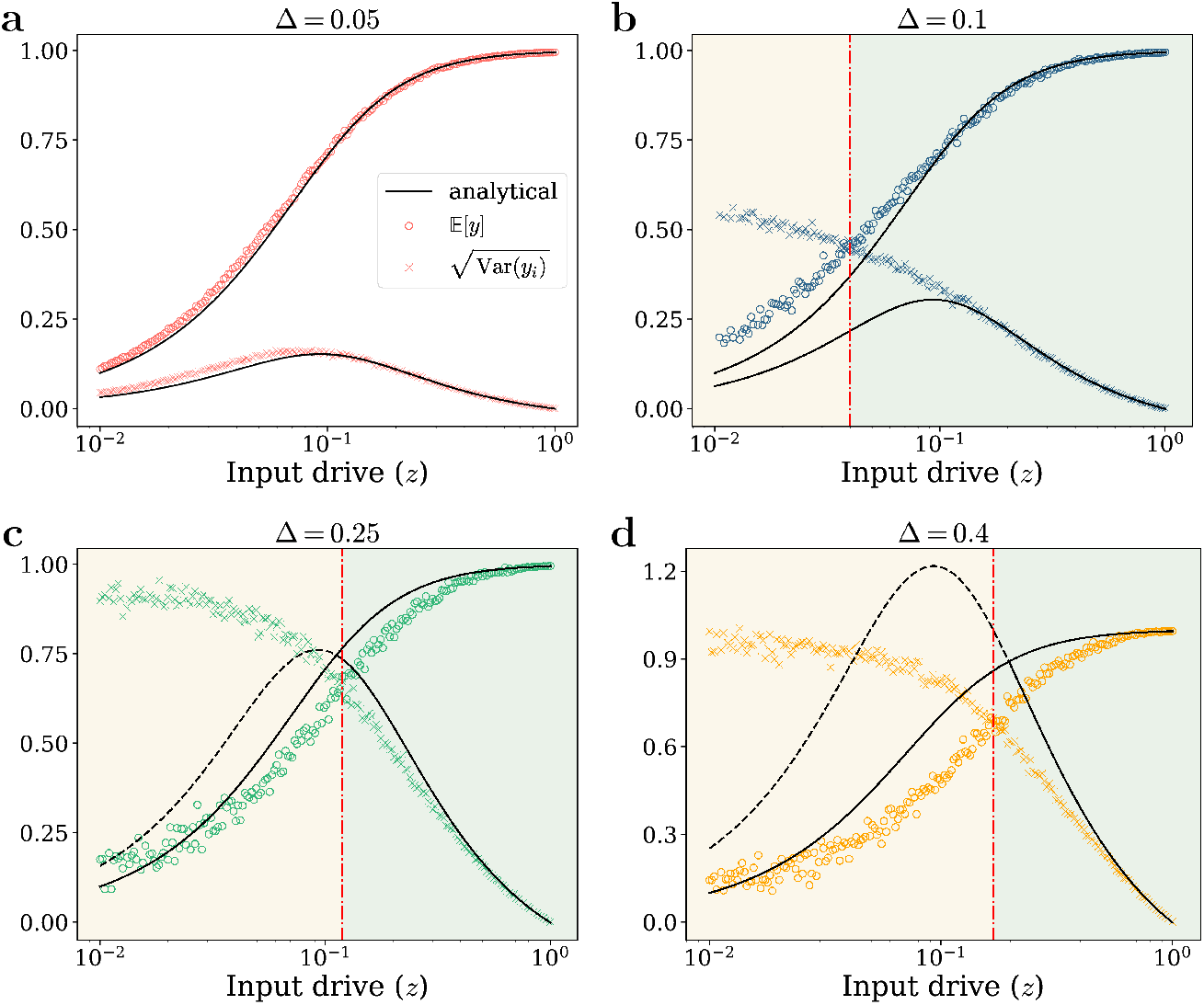
Loss of normalization for <u>localized input drive</u>. **a**, Mean (circles) and standard deviation (crosses) of the fixed point response *y*_*pop*_, defined in Eq. (32), as a function of the norm *z* of the localized input drive *z*_*i*_ = *zδ*_1*i*_ for an E:I balanced recurrence matrix *K* with zero mean and recurrent interaction strength Δ = 0.05. We used *N* = 100 neurons and averaged over 10^3^ realizations of the random matrix *K*. The analytical approximations (solid curves) for the mean Eq. (33) and standard deviation Eq. (34) of the response, computed with perturbation theory, show a good agreement between the theoretical and numerical solutions. In this case the standard deviation is smaller than the mean for all values of *z*, so the neural responses are always normalized. **b, c, d** Same as in **a**, but using Δ = 0.1, 0.25, 0.4. The standard deviation is smaller than the mean at large input drive, but gets bigger than the mean at small input drive. The value of *z* where the two curves cross each other, given by Eq. (37), defines the point at which the neural responses lose normalization. The analytical approximations are in good agreement with numerical simulations for almost all values of the input drive (notice the log scale on the abscissa), but become less accurate at small *z* where the responses are non-normalized.

Since the fixed point *y*_*i*_ depends on *i* we consider the sum over all responses *y*_*pop*_ defined as

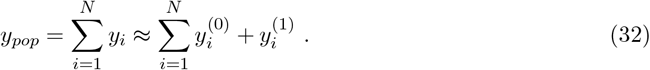

The mean of *y*_*pop*_ is simply

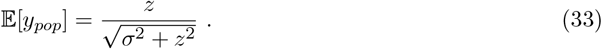

**FIG. S8.**
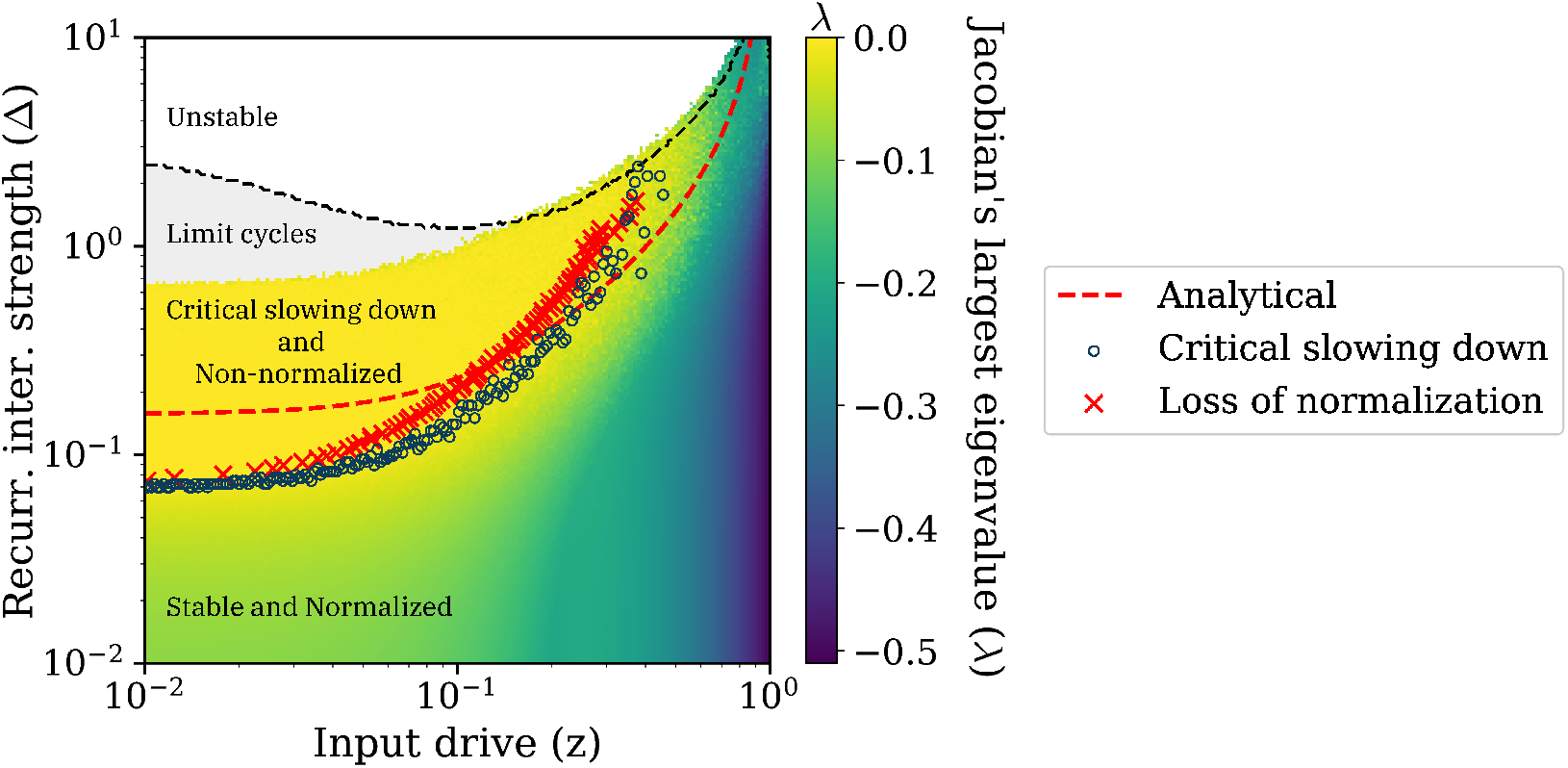
Loss of normalization predicts the onset of critical slowing down for <u>localized input drive</u>. Real part of the largest eigenvalue of the Jacobian at the fixed point in the (*z*, Δ) plane obtained by solving numerically Eqs. (3) with a localized input drive, i.e., *z*_*i*_ = *zδ*_*i*1_. The parameters of ORGaNICs are the same as those used for generating Fig. 5 in the main text. Color represents the maximum value of *λ* across 100 trials. Circuits with small Δ are stable at any value of the input drive *z* and converge quickly to their fixed point, as indicated by a strictly negative eigenvalue *λ <* 0. Conversely, for Δ_*csd*_ *<* Δ *<* Δ_*c*_, the circuits exhibit critical slowing down, in that they approach the fixed point very slowly, as indicated by the zero eigenvalue, *λ* = 0. The onset of critical slowing down is defined by the first time the eigenvalue becomes zero, here denoted by the blue empty circle. The onset of slowing down is equally well captured by the red crosses, representing the boundary between the normalized and non-normalized phases. Loss of normalization is a good proxy for critical slowing down even for localized input drives. For sufficiently large Δ the circuits exhibit limit cycles and for even larger Δ they eventually become unstable, where instability is defined as trajectories diverging in at least 50% of trials.

The variance is given by

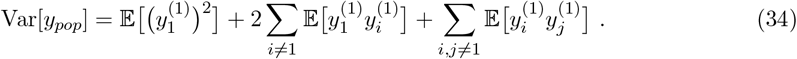

The calculation of the expectation values gives

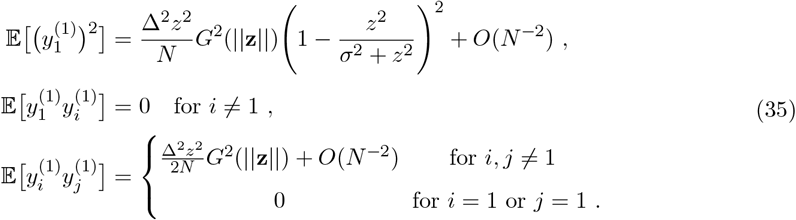

Inserting the previous expressions in Eq. (34) and keeping only the leading order in *N* we find

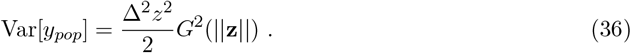

Equating the mean and standard deviation of *y*_*pop*_ we find the Δ_*loss*_(*z*) as

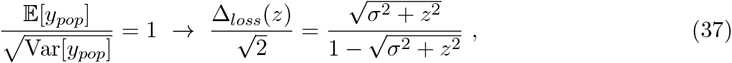

which is the same expression as in Eq. (30).

### D. Frequency of oscillations

To understand how the interplay between external stimuli and recurrent drive affects the dynamics of ORGaNICs, we investigated the system’s propensity to oscillate under varying conditions. Specifically, we explored the influence of the overall input drive *z* and the recurrent interaction strength Δ. We systematically varied these two parameters and computed the average oscillation frequency of the network activity as the mean imaginary part of the Jacobian eigenvalues Im(*λ*_**J**_)*/*(2*π*) evaluated at the system’s fixed point. Oscillatory dynamics (spiralling fixed points) are indicated by complex eigenvalues. Fig. S9 shows the resulting heat map in the (*z*, Δ) plane, where we plot the mean oscillation frequency for a network of *N* = 100 neurons with time constants *τ*_*y*_ = *τ*_*a*_ = 2 msec, considering both delocalized (Fig. S9a) and localized input drives (Fig. S9b).

We find distinct dynamical regimes. When the input drive and the synaptic strength are weak, i.e., *z* ≲ 0.1 and Δ ≲ 0.1, the circuits settle into a stable, non-oscillating fixed point. However, as the input drive increases (*z* ≳ 0.1), the fixed point becomes a spiral attractor, leading to oscillations falling within the gamma frequency range (30-100 Hz). Within this regime, the frequency of these input-driven oscillations scales positively with the input drive *z* (Fig. S9d). A different scenario unfolds when the recurrent synaptic strength is increased (Fig. S9c). For low input drive (*z* ≲ 0.1), damped oscillations emerge beyond Δ ≈ 0.1. In this recurrence-driven regime, the oscillation frequency increases monotonically with Δ from 0 Hz to 80 Hz, before the attractor ultimately turns into limit cycles at higher Δ. These findings highlight the dual roles of external input and internal recurrent drive in shaping the frequency of the oscillatory behavior in ORGaNICs.

**FIG. S9.**
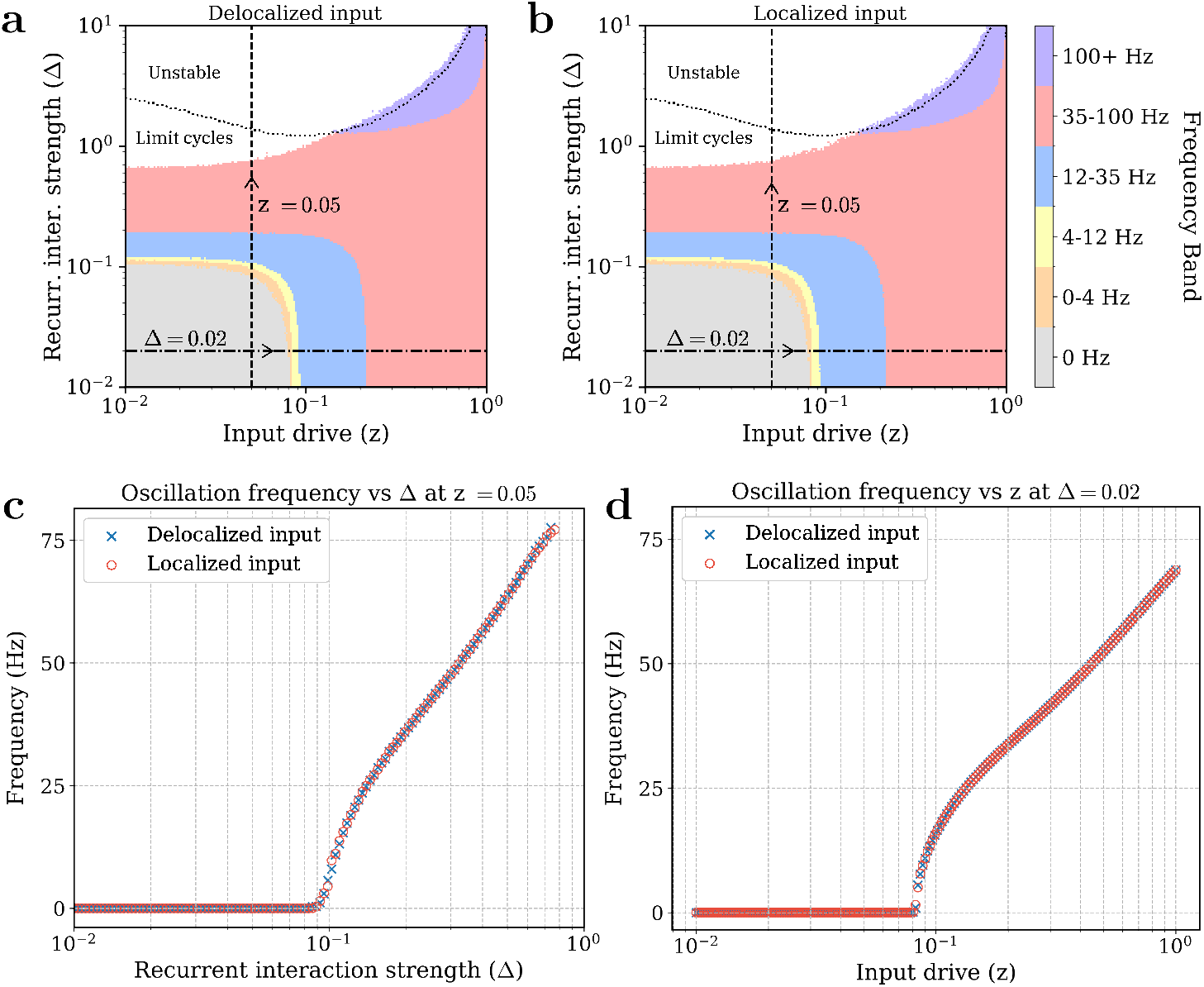
Phase diagram and oscillation frequencies in ORGaNICs. **a, b**, Phase diagrams depicting the average oscillation frequency for ORGaNICs as a function of input drive (*z*) and recurrent interaction strength (Δ). Frequency is color-coded according to standard bands (see color bar: 0 Hz, 0-4 Hz, 4-12 Hz, 12-35 Hz, 35-100 Hz, 100+ Hz), calculated as the mean imaginary part of the Jacobian eigenvalues Im(*λ*_**J**_)*/*(2*π*) across trials. Results are shown for delocalized (**a**) and localized (**b**) inputs in a system with *N* = 100 neurons, semisaturation constant *σ* = 0.1, and time constants *τ*_*y*_ = *τ*_*a*_ = 2 msec. Dotted curves indicate the transition from limit cycles to an unstable regime, where instability is defined as trajectories diverging in at least 50% of trials. **c**, Oscillation frequency vs recurrent interaction strength (Δ) at a fixed input drive *z* = 0.05. **d**, Oscillation frequency vs input drive (*z*) at a fixed recurrent interaction strength Δ = 0.02. Plots **c** and **d** compare delocalized (blue circles) and localized (red circles) inputs, showing minimal difference between the two input types.

### E. ORGaNICs with alternative activation functions

We consider the effect of changing the activation function, which determines the firing rates (*y*^+^) from the membrane potentials (*y*) of the principal neurons. In the model studied in the main text we use a quadratic activation function *y*^+^ = *y*^2^ (see phase diagram of stability in Fig. S10a). Here, we investigate two alternative models incorporating rectification in the activation function, defined as ⌊*x*⌋ = max(0, *x*), a common choice known as ReLU in artificial neural networks.

**FIG. S10.**
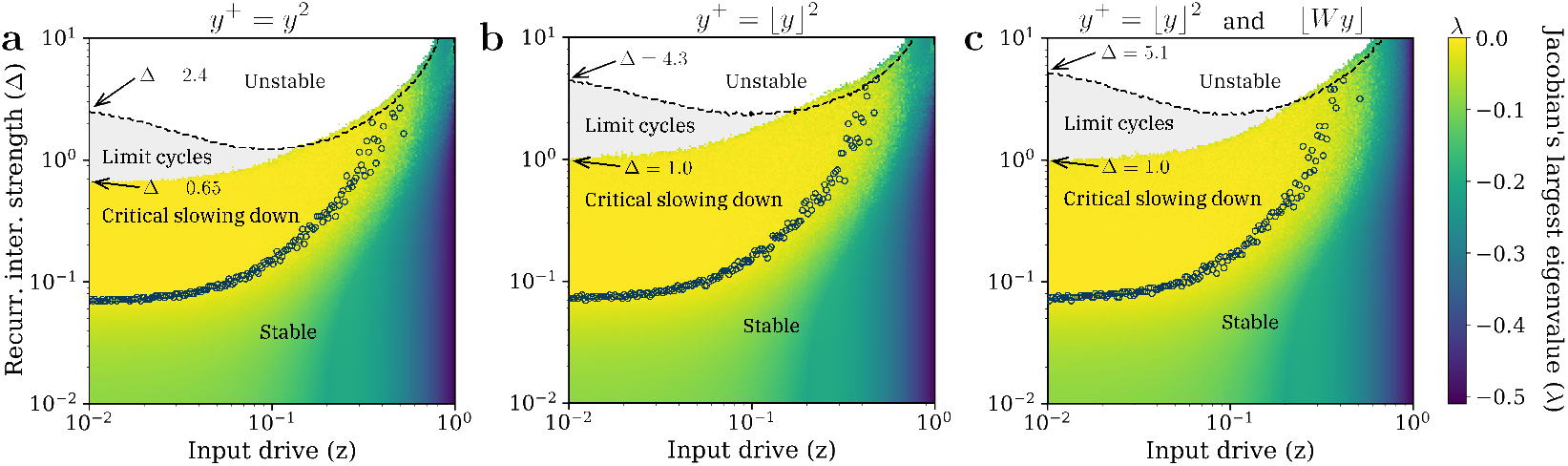
Effect of rectification on network stability. Phase diagram in the parameter space of input drive *z* and recurrent interaction strength Δ (mesh size 200 *×* 200). Color represents the maximum value (across 100 trials) of the Jacobian’s eigenvalue with the largest real part (*λ*). The input type and the parameters of ORGaNICs are the same as those used for generating Fig. 5 in the main text. The panels correspond to: **a**, Model with quadratic activation (*y*^+^ = *y*^2^), identical to the phase diagram in Fig. 5. **b**, Model with rectification in the activation function (*y*^+^ = *⌊y⌋*^2^, Eq. 38). **c**, Model with rectification applied in both the activation function and after the recurrent summation (*y*^+^ = *⌊y⌋*^2^ and *⌊Wy⌋* term, Eq. 39). We observe that the boundaries marking the transition from critical slowing down to limit cycles, and from limit cycles to unstable dynamics (indicated by dashed curves), shift upwards in going from **a** to **b** to **c**.

First, we introduce rectification such that the firing rate is calculated as *y*^+^ = ⌊*y*⌋^2^. This gives us the following dynamical system (see phase diagram in Fig. S10b):

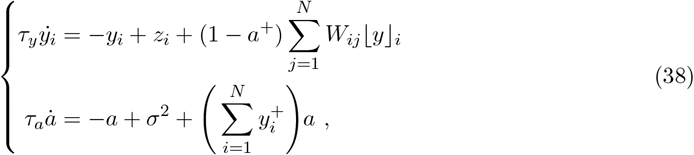

In the second model, we explore a different placement for rectification. While still using the rectified firing rate *y*^+^ = ⌊*y*⌋^2^, we apply rectification after the weighted recurrent inputs have been summed, in the dynamical equation for *y*. This leads to the following dynamical system (see corresponding phase diagram in Fig. S10c):

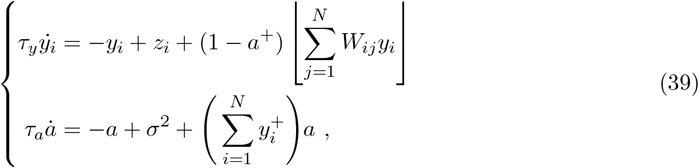

The model in Eq. (39) is not neurobiologically relevant, but is relevant for designing ML architectures [18].

We analyzed the stability of these models by examining their phase diagrams in the parameter space of input drive (*z*) and recurrent interaction strength (Δ), shown in Fig. S10. We find that introducing these alternative forms of rectification do not qualitatively change the network’s phase diagram. However, the boundaries that mark the transition from critical slowing down to limit cycles, and from limit cycles to unstable dynamics, shift towards larger values of Δ, when going from panel (a) to (b) to (c) of Fig. S10. Therefore, introducing rectification increases the range of parameters for which the neuron’s trajectories remain bounded (including the limit cycles regime).

### F. E-I imbalanced recurrent networks

In this section, we investigate the impact of excitation-inhibition (E-I) imbalance in the recurrent weight matrix *W* on the stability of ORGaNICs. We introduce E-I imbalance by setting a non-zero mean *µ* for the entries of the recurrent connectivity matrix *K*, such that *K*_*ii*_ ∼ 𝒩 (*µ/N*, Δ^2^*/N*) and *K*_*ij*_ ∼ 𝒩 (*µ/N*, Δ^2^*/*2*N*) for *i*≠ *j*. This introduces net inhibition (for *µ <* 0) or net excitation (*µ >* 0) in *K*. We generated phase diagrams, shown in Fig. S11, analogous to Fig. 5 for different values of *µ* = [0.0, 0.05, 0.1, 0.25, 0.5, 1.0, −0.1, −0.5, −1.0], using a delocalized input drive 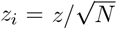 and network parameters *N* = 100, *σ* = 0.1, *τ*_*y*_ = *τ*_*a*_. We observe three main things:

1. increasing excitation (larger positive *µ*) shifts the onset of critical slowing down towards larger values of the recurrent interaction strength Δ. For strong excitatory imbalance (e.g., *µ* = 1.0), the network transitions from the stable regime to the limit cycle regime at small input drive *z* without undergoing critical slowing down;
2. increasing inhibition (larger negative *µ*) makes the circuit operate in the critically slowed down regime for larger values of Δ at any input drive *z*. For strong inhibitory imbalance (e.g., *µ* = −1.0), the circuit remains stable across all *z* without entering into limit cycles at small *z*;
3. most importantly, the loss of normalization is still a good predictor of the onset of critical slowing down across all values of *µ*.

We also examined how E-I imbalance affects the oscillation frequencies in ORGaNICs (Fig. S12). Upon increasing excitation (*µ >* 0), the region exhibiting high-frequency oscillations (gamma band and higher) expands. For strong excitation (e.g., *µ* = 1.0), the network tends to oscillate at high frequencies across a wider range of Δ and *z* values. This suggests that net excitation in the recurrent connections promotes faster oscillations. On the contrary, increasing inhibition (Fig. S12g,h,i) (*µ <* 0) promotes slower oscillations, especially at large input drives.

**FIG. S11.**
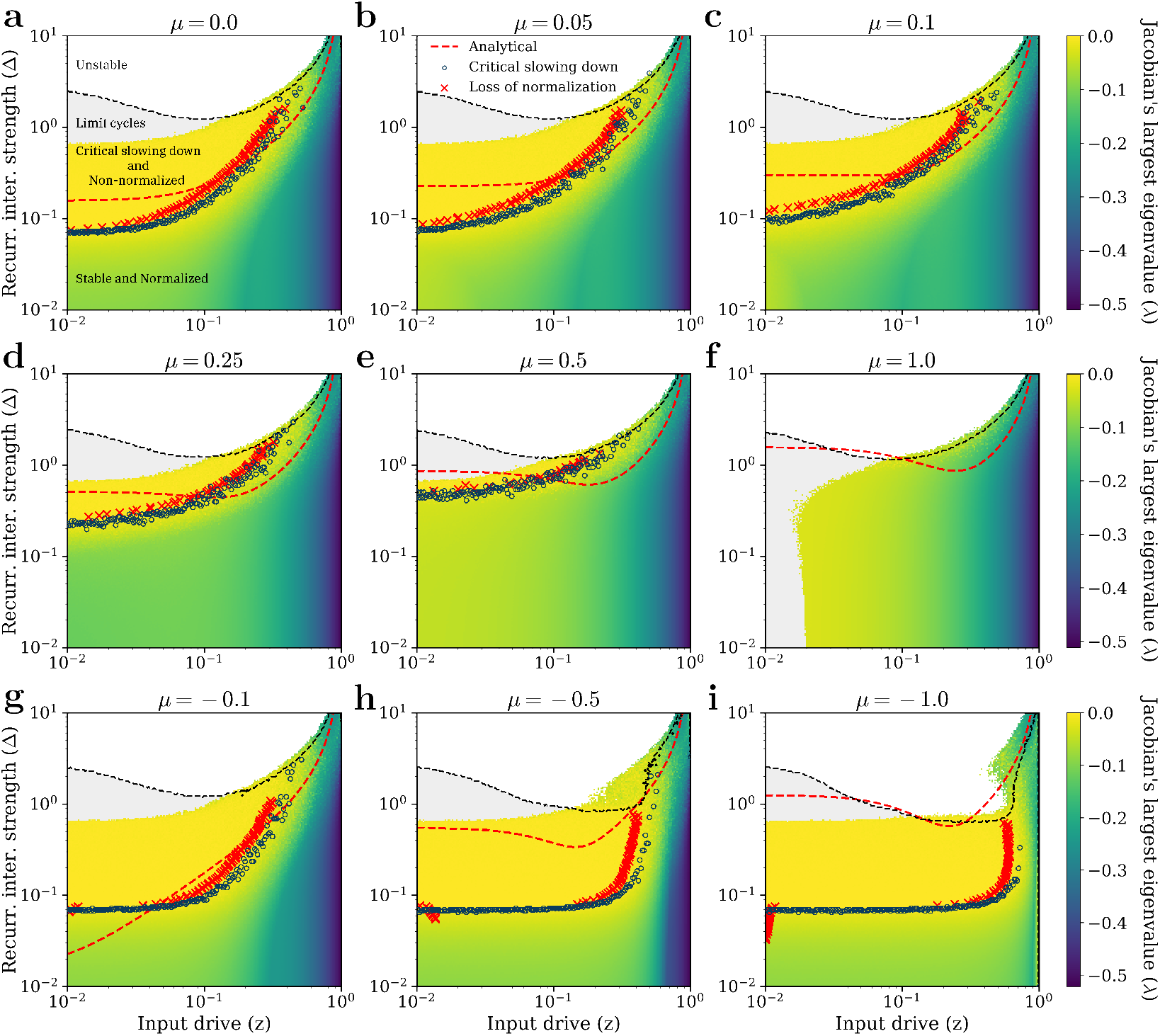
Effect of E-I imbalanced recurrence on stability. Phase diagrams showing the real part of the largest eigenvalue *λ* of the Jacobian at the fixed point in the (*z*, Δ) plane for varying levels of E-I imbalance, controlled by the mean *µ* of the recurrent weights *K*_*ij*_. Each panel corresponds to a different value of *µ* (see discussion in the text): **a**, *µ* = 0.0 (balanced, identical to Fig. 5); excess excitation: **b**, *µ* = 0.05, **c**, *µ* = 0.1, **d**, *µ* = 0.25, **e**, *µ* = 0.5, **f**, *µ* = 1.0; excess inhibition: **g**, *µ* = −0.1, **h**, *µ* = −0.5, **i**, *µ* = −1.0. Color represents the maximum *λ* across 100 trials (for *N* = 100, *σ* = 0.1, *τ*_*y*_ = *τ*_*a*_, delocalized input 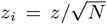). Blue open circles mark the numerically determined onset of critical slowing down, while red crosses indicate the numerically determined loss of normalization. The dashed red curves show the analytical prediction for loss of normalization. Dashed black curves delineate boundaries between the limit cycle (gray region) and the unstable (white region) regimes, where instability is defined as trajectories diverging in at least 50% of trials.

**FIG. S12.**
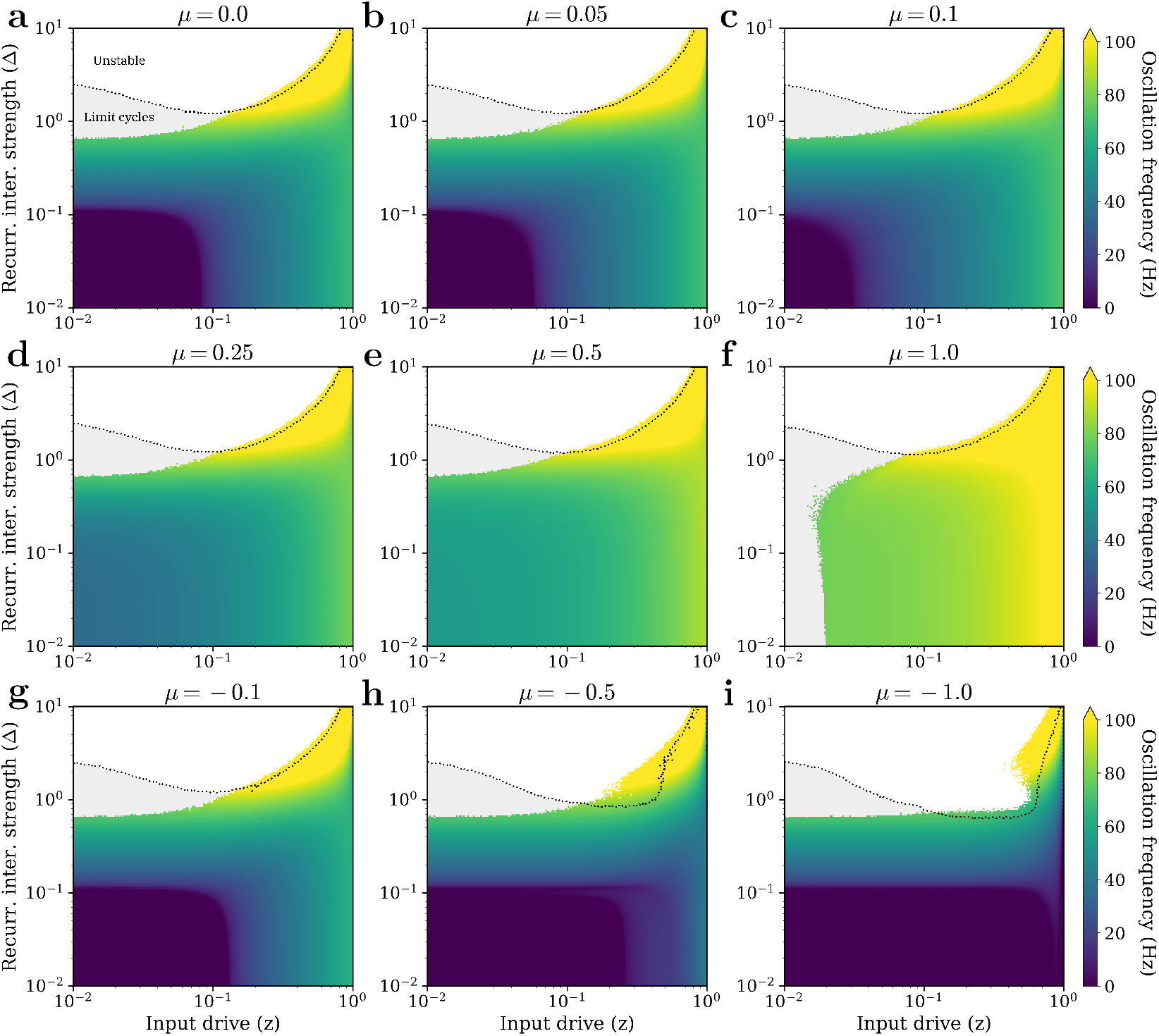
Effect of E-I imbalanced recurrence on oscillation frequency. Phase diagrams depicting the average oscillation frequency (calculated as mean of Im(*λ*_**J**_)*/*(2*π*) across trials) as a function of input drive *z* and recurrent interaction strength Δ for varying levels of E-I imbalance *µ*. Parameters are *N* = 100, *σ* = 0.1, *τ*_*y*_ = *τ*_*a*_ = 2 msec, delocalized input 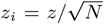. Panels correspond to: **a**, *µ* = 0.0; excess excitation: **b**, *µ* = 0.05, **c**, *µ* = 0.1, **d**, *µ* = 0.25, **e**, *µ* = 0.5, **f**, *µ* = 1.0; excess inhibition: **g**, *µ* = −0.1, **h**, *µ* = −0.5, **i**, *µ* = −1.0. Dotted black curves delineate boundaries between the limit cycle (gray region) and the unstable (white region) regimes, where instability is defined as trajectories diverging in at least 50% of trials. Overall, increasing excitation (*µ >* 0) generally promotes higher oscillation frequencies, especially at small input drives. While increasing inhibition (*µ <* 0) promotes lower oscillation frequencies, especially at large input drives.

### G. Effect of critical slowing down on neural variability

A key marker of critical slowing down is a drastic increase in trial-to-trial neural variability and noise correlations between neurons. To illustrate why this occurs, we consider the ORGaNICs model with additive Gaussian white noise in the dynamical system:

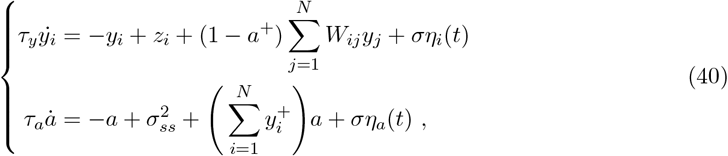

where *η*_*i*_(*t*) and *η*_*a*_(*t*) represent uncorrelated Gaussian white noise processes with zero mean and unit variance (i.e., 𝔼[*η*_*k*_(*t*)*η*_*l*_(*s*)] = *δ*_*kl*_*δ*(*t* − *s*)), and *σ* here denotes the strength of this noise. Note that 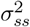 is used for the semisaturation constant to avoid confusion with the noise strength *σ*. This introduction of stochasticity is distinct from the randomness in the recurrent matrix *W* considered in the main body of the manuscript.

Assuming the dynamical system operates in the vicinity of the stable fixed point (found in both stable and critically slowed-down regimes) and that the noise strength *σ* is sufficiently small, we can linearize the system around the fixed point. Let **x** be the vector of deviations from the fixed point. The linearized system is:

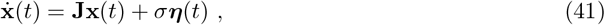

where **J** is the Jacobian matrix evaluated at the fixed point, and ***η***(*t*) is the vector of white noise processes. The steady-state covariance matrix **P** = 𝔼[**xx**^⊤^] (whose diagonal entries capture trial-to-trial variability and off-diagonal entries capture noise correlations) is given by the solution to the continuous-time Lyapunov equation:

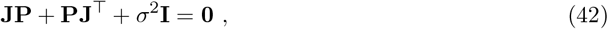

where **I** is the identity matrix, and *σ*^2^**I** is the covariance matrix of the noise term *σ****η***(*t*).

Assuming that **J** is diagonalizable, we can write its eigendecomposition as **J** = **VΛV**^−1^, where **Λ** is a diagonal matrix whose entries *λ*_*i*_ are the eigenvalues of **J**, and **V** is the matrix whose columns are the corresponding eigenvectors. We can transform the coordinates to the eigenbasis of **J** by defining **y** = **V**^−1^**x**. The covariance matrix of **y** is **M** = 𝔼[**yy**^⊤^] = **V**^−1^**P**(**V**^−1^)^⊤^ = **V**^−1^**PV**^−⊤^. Left-multiplying Eq. (42) by **V**^−1^ and right-multiplying by **V**^−⊤^, we obtain:

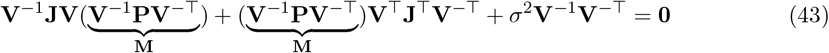

Using **V**^−1^**JV** = **V**^⊤^**J**^⊤^**V**^−⊤^ = **Λ** and defining **B** = **V**^−1^**V**^−⊤^, Eq. (43) becomes:

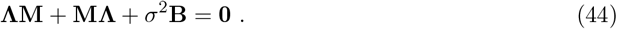

This is the Lyapunov equation for the transformed coordinates **y**. Since **Λ** is diagonal, we can solve for the entries of **M** element-wise:

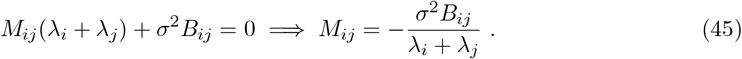

Therefore, if the real part of an eigenvalue, say Re(*λ*_*k*_), approaches zero (which characterizes critical slowing down), the denominator 2Re(*λ*_*k*_) for the diagonal term *M*_*kk*_ becomes very small. Assuming *B*_*kk*_ (which depends on the eigenvectors) is non-zero, *M*_*kk*_ will become very large:

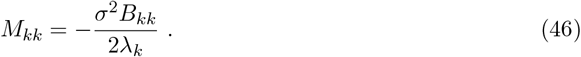

As Re(*λ*_*k*_) → 0, the magnitude of *M*_*kk*_ tends to infinity. Since the original covariance matrix entries *P*_*mn*_ are linear combinations of *M*_*ij*_ (as **P** = **VMV**^⊤^), a large *M*_*kk*_ will typically lead to large entries *P*_*mn*_. This implies increased trial-to-trial variability (large diagonal elements of **P**) and large noise correlations (large off-diagonal elements of **P**) when the system is in the critically slowed-down regime compared to the stable and normalized regime.

## REFERENCES

[1] David J. Heeger. Normalization of cell responses in cat striate cortex. Visual Neuroscience, 9(2): 181–197, 1992. doi:10.1017/S0952523800009640.

[2] Matteo Carandini and David J. Heeger. Summation and division by neurons in primate visual cortex. Science, 264(5163):1333–1336, 1994. doi:10.1126/science.8191289. URL https://www.science.org/doi/abs/10.1126/science.8191289.

[3] Shawn R Olsen, Vikas Bhandawat, and Rachel I Wilson. Divisive normalization in olfactory population codes. Neuron, 66(2):287–299, 2010.

[4] Odelia Schwartz and Eero Simoncelli. Natural sound statistics and divisive normalization in the auditory system. Advances in neural information processing systems, 13, 2000.

[5] Matteo Carandini and David J Heeger. Normalization as a canonical neural computation. Nature reviews neuroscience, 13(1):51–62, 2012.

[6] John H Reynolds and David J Heeger. The normalization model of attention. Neuron, 61(2):168–185, 2009.

[7] Wei Ji Ma, Masud Husain, and Paul M Bays. Changing concepts of working memory. Nature neuro-science, 17(3):347–356, 2014.

[8] Kenway Louie, Lauren E Grattan, and Paul W Glimcher. Reward value-based gain control: divisive normalization in parietal cortex. Journal of Neuroscience, 31(29):10627–10639, 2011.

[9] Frances S Chance, L.F Abbott, and Alex D Reyes. Gain modulation from background synaptic input. Neuron, 35(4):773–782, 2002. ISSN 0896-6273. doi:10.1016/S0896-6273(02)00820-6. URL https://www.sciencedirect.com/science/article/pii/S0896627302008206.

[10] Matteo Carandini, David J Heeger, and Walter Senn. A synaptic explanation of suppression in visual cortex. Journal of Neuroscience, 22(22):10053–10065, 2002. ISSN 0270-6474. doi:10.1523/JNEUROSCI.22-22-10053.2002. URL https://www.jneurosci.org/content/22/22/10053.

[11] Hirofumi Ozeki, Ian M Finn, Evan S Schaffer, Kenneth D Miller, and David Ferster. Inhibitory stabilization of the cortical network underlies visual surround suppression. Neuron, 62(4):578–592, 2009.

[12] Hillel Adesnik and Massimo Scanziani. Lateral competition for cortical space by layer-specific horizontal circuits. Nature, 464(7292):1155–1160, 2010.

[13] Tatsuo K Sato, Bilal Haider, Michael Häusser, and Matteo Carandini. An excitatory basis for divisive normalization in visual cortex. Nature neuroscience, 19(4):568–570, 2016.

[14] Hillel Adesnik. Synaptic mechanisms of feature coding in the visual cortex of awake mice. Neuron, 95 (5):1147–1159, 2017.

[15] Kevin A Bolding and Kevin M Franks. Recurrent cortical circuits implement concentration-invariant odor coding. Science, 361(6407):eaat6904, 2018.

[16] David J. Heeger and Wayne E. Mackey. Oscillatory recurrent gated neural integrator circuits (organics), a unifying theoretical framework for neural dynamics. Proceedings of the National Academy of Sciences, 116(45):22783–22794, 2019. doi:10.1073/pnas.1911633116. URL https://www.pnas.org/doi/abs/10.1073/pnas.1911633116.

[17] S.C. Cannon, D.A. Robinson, and S. Shamma. A proposed neural network for the integrator of the oculomotor system. Biol. Cybernetics, 49:127–136, 1983. doi:10.1007/BF00320393. URL https://www.sciencedirect.com/science/article/pii/S0896627302008206.

[18] Shivang Rawat, David Heeger, and Stefano Martiniani. Unconditional stability of a recurrent neural circuit implementing divisive normalization. Advances in Neural Information Processing Systems, 37: 14712–14750, 2024.

[19] EP Wigner. Statistical properties of real symmetric matrices with many dimensions can. Math. Congr. Proc., University of Toronto Press, Toronto, page 174, 1957.

[20] Lei Dai, Daan Vorselen, Kirill S Korolev, and Jeff Gore. Generic indicators for loss of resilience before a tipping point leading to population collapse. Science, 336(6085):1175–1177, 2012.

[21] Marten Scheffer, Stephen R Carpenter, Timothy M Lenton, Jordi Bascompte, William Brock, Vasilis Dakos, Johan Van de Koppel, Ingrid A Van de Leemput, Simon A Levin, Egbert H Van Nes, et al. Anticipating critical transitions. science, 338(6105):344–348, 2012.

[22] Wilson S Geisler and Duane G Albrecht. Visual cortex neurons in monkeys and cats: detection, discrimination, and identification. Visual neuroscience, 14(5):897–919, 1997.

[23] AB Bonds. Role of inhibition in the specification of orientation selectivity of cells in the cat striate cortex. Visual neuroscience, 2(1):41–55, 1989.

[24] GC DeAngelis, JG Robson, I Ohzawa, and RD Freeman. Organization of suppression in receptive fields of neurons in cat visual cortex. Journal of Neurophysiology, 68(1):144–163, 1992.

[25] Gregory C DeAngelis, RALPH D Freeman, and IZUMI Ohzawa. Length and width tuning of neurons in the cat’s primary visual cortex. Journal of neurophysiology, 71(1):347–374, 1994.

[26] Wyeth Bair, James R Cavanaugh, and J Anthony Movshon. Time course and time-distance relation-ships for surround suppression in macaque v1 neurons. Journal of Neuroscience, 23(20):7690–7701, 2003.

[27] Nicole C Rust, Valerio Mante, Eero P Simoncelli, and J Anthony Movshon. How mt cells analyze the motion of visual patterns. Nature neuroscience, 9(11):1421–1431, 2006.

[28] James R Cavanaugh, Wyeth Bair, and J Anthony Movshon. Nature and interaction of signals from the receptive field center and surround in macaque v1 neurons. Journal of neurophysiology, 88(5): 2530–2546, 2002.

[29] James R Cavanaugh, Wyeth Bair, and J Anthony Movshon. Selectivity and spatial distribution of signals from the receptive field surround in macaque v1 neurons. Journal of neurophysiology, 88(5): 2547–2556, 2002.

[30] Edward H Adelson and James R Bergen. Spatiotemporal energy models for the perception of motion. Journal of the optical society of america A, 2(2):284–299, 1985.

[31] Matteo Carandini. Amplification of trial-to-trial response variability by neurons in visual cortex. PLoS biology, 2(9):e264, 2004.

[32] Jeffrey S Anderson, Ilan Lampl, Deda C Gillespie, and David Ferster. The contribution of noise to contrast invariance of orientation tuning in cat visual cortex. Science, 290(5498):1968–1972, 2000.

[33] Nicholas J Priebe and David Ferster. Inhibition, spike threshold, and stimulus selectivity in primary visual cortex. Neuron, 57(4):482–497, 2008.

[34] Katie A Ferguson and Jessica A Cardin. Mechanisms underlying gain modulation in the cortex. Nature Reviews Neuroscience, 21(2):80–92, 2020.

[35] Brendan K Murphy and Kenneth D Miller. Multiplicative gain changes are induced by excitation or inhibition alone. Journal of neuroscience, 23(31):10040–10051, 2003.

[36] Hugh R Wilson. Binocular contrast, stereopsis, and rivalry: Toward a dynamical synthesis. Vision research, 140:89–95, 2017.

[37] Jian Ding and George Sperling. A gain-control theory of binocular combination. Proceedings of the National Academy of Sciences, 103(4):1141–1146, 2006.

[38] Tim S Meese, Mark A Georgeson, and Daniel H Baker. Binocular contrast vision at and above threshold. Journal of vision, 6(11):7–7, 2006.

[39] Kenneth D Miller and Todd W Troyer. Neural noise can explain expansive, power-law nonlinearities in neural response functions. Journal of neurophysiology, 87(2):653–659, 2002.

[40] Larry F Abbott, JA Varela, Kamal Sen, and SB Nelson. Synaptic depression and cortical gain control. Science, 275(5297):221–224, 1997.

[41] R. M. May. Will a large complex system be stable? Nature, 238:413–414, 1972. doi: 10.1038/238413a0.

[42] Stefano Allesina and Si Tang. Stability criteria for complex ecosystems. Nature, 483(7388):205–208, 2012.

[43] Flaviano Morone, Gino Del Ferraro, and Hernán A Makse. The k-core as a predictor of structural collapse in mutualistic ecosystems. Nature physics, 15(1):95–102, 2019.

[44] K. S. McCann. The diversity–stability debate. Nature, 405:228–233, 2000. doi: 10.1038/35012234.

[45] Francesca Arese Lucini, Flaviano Morone, Maria Silvina Tomassone, and Hernán A Makse. Diversity increases the stability of ecosystems. PloS one, 15(4):e0228692, 2020.

[46] Ian A Hatton, Onofrio Mazzarisi, Ada Altieri, and Matteo Smerlak. Diversity begets stability: Sublinear growth and competitive coexistence across ecosystems. Science, 383(6688):eadg8488, 2024.

[47] C. van Vreeswijk and H. Sompolinsky. Chaos in neuronal networks with balanced excitatory and inhibitory activity. Science, 274(5293):1724–1726, 1996. doi:10.1126/science.274.5293.1724. URL https://www.science.org/doi/abs/10.1126/science.274.5293.1724.

[48] Robbe LT Goris, Ruben Coen-Cagli, Kenneth D Miller, Nicholas J Priebe, and Máté Lengyel. Response sub-additivity and variability quenching in visual cortex. Nature Reviews Neuroscience, 25(4):237–252, 2024.

[49] Shivang Rawat and Stefano Martiniani. Element-wise and recursive solutions for the power spectral density of biological stochastic dynamical systems at fixed points. Physical Review Research, 6(4): 043179, 2024.

[50] Marten Scheffer, Jordi Bascompte, William A Brock, Victor Brovkin, Stephen R Carpenter, Vasilis Dakos, Hermann Held, Egbert H Van Nes, Max Rietkerk, and George Sugihara. Early-warning signals for critical transitions. Nature, 461(7260):53–59, 2009.

[51] Matias I Maturana, Christian Meisel, Katrina Dell, Philippa J Karoly, Wendyl D’Souza, David B Grayden, Anthony N Burkitt, Premysl Jiruska, Jan Kudlacek, Jaroslav Hlinka, et al. Critical slowing down as a biomarker for seizure susceptibility. Nature communications, 11(1):2172, 2020.

[52] URI Polat, DOV Sagi, and Anthony M Norcia. Abnormal long-range spatial interactions in amblyopia. Vision research, 37(6):737–744, 1997.

[53] Dave Ellemberg, Robert F Hess, and A Serge Arsenault. Lateral interactions in amblyopia. Vision Research, 42(21):2471–2478, 2002.

[54] Vittorio Porciatti, Paolo Bonanni, Adriana Fiorentini, and Renzo Guerrini. Lack of cortical contrast gain control in human photosensitive epilepsy. Nature neuroscience, 3(3):259–263, 2000.

[55] Jeffrey J Tsai, Anthony M Norcia, Justin M Ales, and Alex R Wade. Contrast gain control abnormalities in idiopathic generalized epilepsy. Annals of neurology, 70(4):574–582, 2011.

[56] Jean-Paul Noel and Dora E Angelaki. A theory of autism bridging across levels of description. Trends in Cognitive Sciences, 27(7):631–641, 2023.

[57] Ari Rosenberg, Jaclyn Sky Patterson, and Dora E Angelaki. A computational perspective on autism. Proceedings of the National Academy of Sciences, 112(30):9158–9165, 2015.

[58] Julie D Golomb, Jenika RB McDavitt, Barbara M Ruf, Jason I Chen, Aybala Saricicek, Kathleen H Maloney, Jian Hu, Marvin M Chun, and Zubin Bhagwagar. Enhanced visual motion perception in major depressive disorder. Journal of Neuroscience, 29(28):9072–9077, 2009.

[59] Anita Must, Zoltán Janka, György Benedek, and Szabolcs Kéri. Reduced facilitation effect of collinear flankers on contrastdetection reveals impaired lateral connectivity in the visual cortex of schizophrenia patients. Neuroscience letters, 357(2):131–134, 2004.

[60] Pamela D Butler, Vance Zemon, Isaac Schechter, Alice M Saperstein, Matthew J Hoptman, Kelvin O Lim, Nadine Revheim, Gail Silipo, and Daniel C Javitt. Early-stage visual processing and cortical amplification deficits in schizophrenia. Archives of general psychiatry, 62(5):495–504, 2005.

[61] Steven Dakin, Patricia Carlin, and David Hemsley. Weak suppression of visual context in chronic schizophrenia. Current Biology, 15(20):R822–R824, 2005.

[62] Duje Tadin, Jejoong Kim, Mikisha L Doop, Crystal Gibson, Joseph S Lappin, Randolph Blake, and Sohee Park. Weakened center-surround interactions in visual motion processing in schizophrenia. Journal of Neuroscience, 26(44):11403–11412, 2006.

[63] Iris IA Groen, Giovanni Piantoni, Stephanie Montenegro, Adeen Flinker, Sasha Devore, Orrin Devinsky, Werner Doyle, Patricia Dugan, Daniel Friedman, Nick F Ramsey, et al. Temporal dynamics of neural responses in human visual cortex. Journal of Neuroscience, 42(40):7562–7580, 2022.

[64] Wyeth Bair and J Anthony Movshon. Adaptive temporal integration of motion in direction-selective neurons in macaque visual cortex. Journal of Neuroscience, 24(33):7305–7323, 2004.

[65] Ilan Dinstein, David J Heeger, Lauren Lorenzi, Nancy J Minshew, Rafael Malach, and Marlene Behrmann. Unreliable evoked responses in autism. Neuron, 75(6):981–991, 2012.

[66] Ilan Dinstein, David J Heeger, and Marlene Behrmann. Neural variability: friend or foe? Trends in cognitive sciences, 19(6):322–328, 2015.

[67] Xiaowen Chen and William Bialek. Searching for long timescales without fine tuning. Physical Review E, 110(3):034407, 2024.

[68] Daniel B Rubin, Stephen D Van Hooser, and Kenneth D Miller. The stabilized supralinear network: a unifying circuit motif underlying multi-input integration in sensory cortex. Neuron, 85(2):402–417, 2015.

